# Morphological principles of neuronal mitochondria

**DOI:** 10.1101/2021.03.15.435547

**Authors:** Rachel Mendelsohn, Guadalupe C. Garcia, Thomas M. Bartol, Christopher T. Lee, P. Khandelwal, Emily Liu, Donald J. Spencer, Adam Husar, Eric A. Bushong, Sebastien Phan, Guy Perkins, Mark H. Ellisman, Alexander Skupin, Terrence J. Sejnowski, Padmini Rangamani

**Affiliations:** Computational Neurobiology Laboratory, Salk Institute for Biological Studies, La Jolla, CA 92037; Mechanical and Aerospace Engineering, University of California, San Diego, La Jolla, CA 92093; National Center for Microscopy and Imaging Research, Center for Research in Biological Systems, Department of Neuroscience, University of California, School of Medicine, San Diego, La Jolla, CA 92093; Luxembourg Centre for Systems Biomedicine, University of Luxembourg, Belvaux, L-4367; Division of Biological Sciences, University of California, San Diego, La Jolla, CA 92093

## Abstract

In the highly dynamic metabolic landscape of a neuron, mitochondrial membrane architectures can provide critical insight into the unique energy balance of the cell. Current theoretical calculations of functional outputs like ATP and heat often represent mitochondria as idealized geometries and therefore can miscalculate the metabolic fluxes. To analyze mitochondrial morphology in neurons of mouse cerebellum neuropil, 3D tracings of complete synaptic and axonal mitochondria were constructed using a database of serial TEM tomography images and converted to watertight meshes with minimal distortion of the original microscopy volumes with a granularity of 1.6 nanometer isotropic voxels. The resulting *in silico* representations were subsequently quantified by differential geometry methods in terms of the mean and Gaussian curvatures, surface areas, volumes, and membrane motifs, all of which can alter the metabolic output of the organelle. Finally, we identify structural motifs that are present across this population of mitochondria; observations which may contribute to future modeling studies of mitochondrial physiology and metabolism in neurons.

## Introduction

Significant amounts of energy are required in the brain to maintain the flow of information [17]. Neurons, as the main active cell type participating in these information flows, are highly polarized cells with extended axons, and exhibit fast electrochemical signals based on ion redistribution, which imposes an energetic burden on them [31]. The energy requirements of a synapse where information is transmitted from one neuron to another have been estimated for the different subprocesses at multiple millions of adenosine triphosphate (ATP) molecules per second [2, 17]. Presynaptic mitochondria are known to be strategically located to respond to the local energy demand of synaptic transmission and have specialized features on the molecular and morphological level [9, 40]. Mitochondrial dysfunction and synaptic homeostasis are believed to contribute to neurodegeneration [9, 32]. The connection between neuronal metabolic dysfunction and mitochondrial ultrastructure has been documented for diseases such as Leigh syndrome, where models correlate insufficient ATP production with compromised morphology [43]. Therefore, understanding the interplay between molecular and morphological organization of mitochondria may contribute new insights into brain energy homeostasis, synaptic function, and mechanisms of neurodegeneration.

Mitochondrial architecture consists of two compositionally and functionally distinct membranes: the outer membrane (OM) and the inner membrane (IM), shown in Figure 1A. The IM is composed of two contiguous structures: the inner boundary membrane (IBM) and the cristae membrane (CM). The IBM is the outer surface which remains in close opposition to the OM (Figure 1B), while the CM is the internal network of compartments connected to the IBM by narrow tubular structures known as cristae junctions (CJs) [35, 12]. This compartmentalized structure provides a highly specialized and adaptable framework for the conversion of metabolic substrates to ATP. The matrix, a volume encapsulated by the IBM and CM, consists of the molecular components needed for the tricarboxylic acid cycle (TCA). The CM itself is the site of the electron transport chain (ETC). Further localization of ETC complexes within the CM is facilitated by variations in membrane curvature, which have been shown to correlate with protein placement and function [6]. The morphology of cristae compartments is often described as either lamellar (sheet-like) or tubular, with a single neuronal mitochondrion commonly exhibiting both types [39]. While a tubular region will have relatively uniform curvature, a lamellar region consists of two nearly flat sheets connected by regions of intense curvature. In addition to promoting a non-uniform distribution of proteins, the highly variable surface created by these structures can optimize the internal surface area and is thought to control the reaction rates of ETC enzymes as well as the formation of supercomplexes [6].

**Figure 1:**
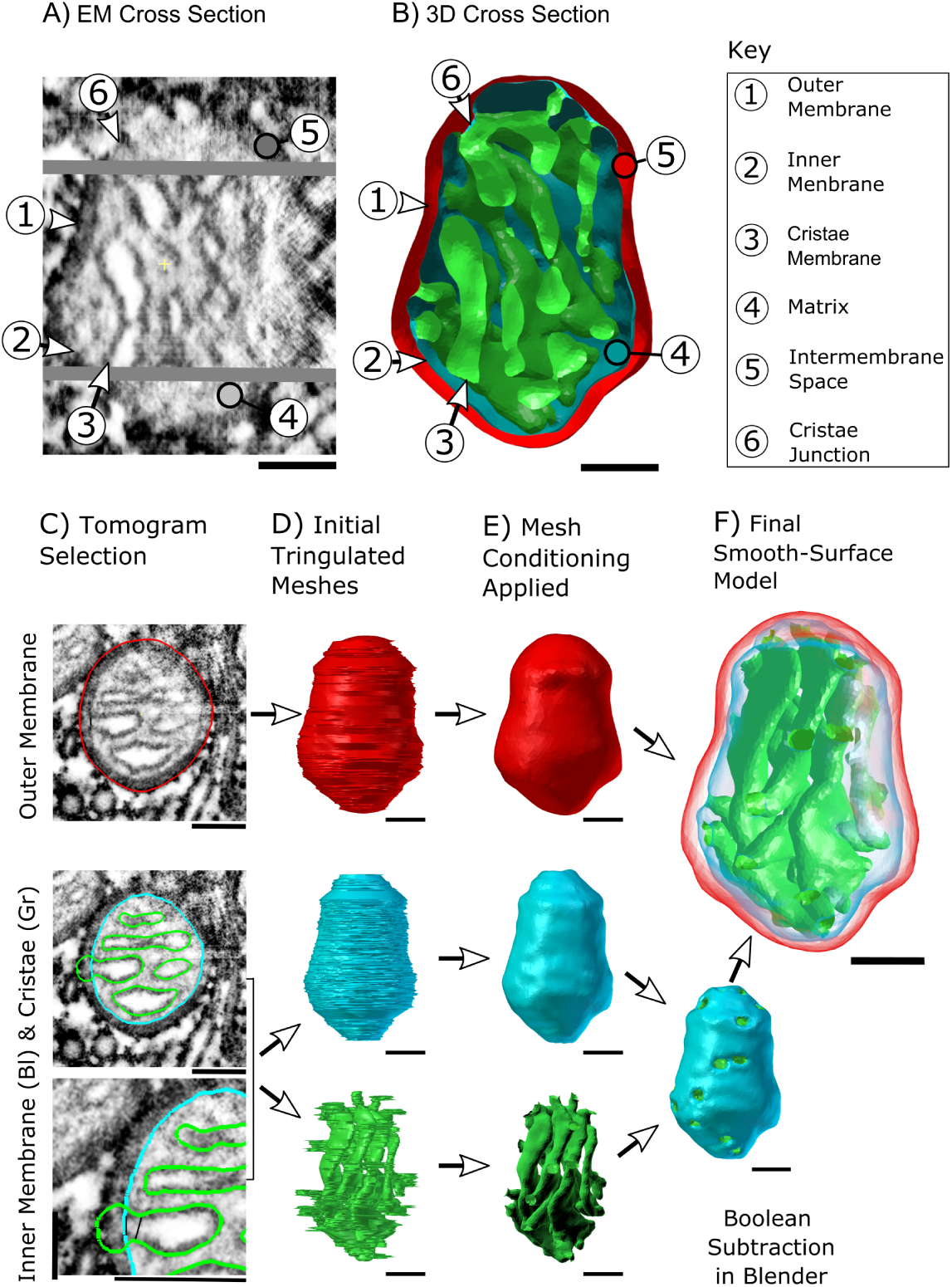
From high resolution 2D representations of mitochondria to high quality 3D reconstructions. (A-B) Serial electron tomographic images of mouse cerebellum neuropil at an isotropic voxel size of 1.64 nm are used to reconstruct the two lipid bilayer membranes of mitochondria. One specific EM cross section is shown here, highlighting the physical section in the tomogram, and its associated 3D cross section reconstruction. The spatial organization of these membranes generates a number of compartments and submembrane regions specified here. (C) The hand-traced contours used in the reconstructions are shown overlayed on a cutout from the respective tomographic image. These three overlays show traces of the three membrane structures; the outer membrane (OM, in red), inner boundary membrane (IBM, in blue), and cristae membrane (CM, in green). Although the inner membrane (IM) is a single structure comprised of the IBM and CM, these two components are treated as separate objects and are combined later. (D) The 3D reconstructions of the membranes are created using the program Contour Tiler [10]. Jaggedness in the model’s surface was found to be due to noise resulting from human tracing error. (E) The noise is smoothed out using GAMer 2 mesh conditioning operations [24] to create the smooth-surface models shown here. The IBM and CM have been joined into one object via Boolean subtraction. (F) All meshes are made transparent and shown together. All Scale Bars: 100 nm.

Within the dense network of tunnels forming the cristae membrane, both low-energy and high-energy structures exist simultaneously. The protein localization and operation which depend on this complex framework make morphometric analysis essential for understanding and modeling mitochondrial function [14]. Recent advances in electron microscopic tomography [41] and reconstruction methods allow for the creation of three-dimensional models of mitochondria with nanometer-level detail [39, 40, 38, 43]. In this work, we systematically and quantitatively reconstruct and analyze the ultrastructural and geometric features of a population of presynaptic mitochondria from mouse cerebellum. Each mitochondrion is reconstructed in its entirety, allowing for the study of mitochondria as functional units. We use our analysis methods to identify the key geometric features of these mitochondria such as surface area-to-volume relationships and surface curvatures, which are essential markers of metabolic activity.

These analyses have been used to identify the structural motifs that define mitochondrial structure, elucidating the physical framework on which presynaptic energy production occurs. To our knowledge, this is the first quantitative morphological analysis of multiple near-complete mitochondria from presynaptic neurons, which includes their surface curvatures and functional motifs. This lays the foundation for defining structure-function relationships in these critical organelles.

## Results

To analyze the architecture of neuronal mitochondria from mouse cerebellum neuropil, 3D reconstructions of 8 presynaptic and 4 axonal mitochondria were generated using serial TEM tomography images (Figure 1A-B). Nine of these organelles were captured at high enough image quality to fully reconstruct the cristae membrane, allowing for complete models of all membranes. From manual membrane traces (Figure 1C) we created high quality meshes (Figure 1D-F) at a scale of 1.6 nm isotropic voxel size. New procedures and tools, including the mesh processing software GAMer 2 [23, 24], state-of-the-art image alignment [57] and gap interpolation techniques, were implemented to minimize distortion from the original microscopy images (see Methods for further details). The fine detail of the resulting smooth meshes (Figure 1E-F) is thus well-preserved for point-by-point calculation of curvature, a comprehensive method for quantifying geometric features of a surface [49, 22]. As can be appreciated in Figure 1E, this level of detail makes it possible to perform such analysis on features as small as cristae junctions which are approximately 25 nm wide [35, 43]. Each of the three membrane components were characterized using these procedures, beginning with the outer membrane.

### Geometric features of the outer membrane reveal a diversity of mitochondrial shapes

Due to their large variability (Figure 2A), we begin by analyzing the geometric features of the outer membrane of 12 mitochondria. For each of these mitochondria, we calculated the volume, surface area, radii along the length, and total length of the outer membrane. These features are represented in Figure 2B–D, and in the supplementary material, a reference number is given in the figures and tables associated to each organelle (Table 4 in the Appendix). The volume enclosed by the OM spans a wide range across the reconstructed mitochondria (Figure 2B), almost by an order of magnitude; the smallest measured volume was (0.0062 μm^3^) while the largest was (0.0459 μm^3^). The surface area of the OM scales with the volume following a power law (Figure 2B, *S* = *αV^γ^*) with coefficient *γ* of 0.74 and *α* of 7.71 (with *R^2^* of 0.97 and p-value of 1 × 10^−8^).

**Figure 2:**
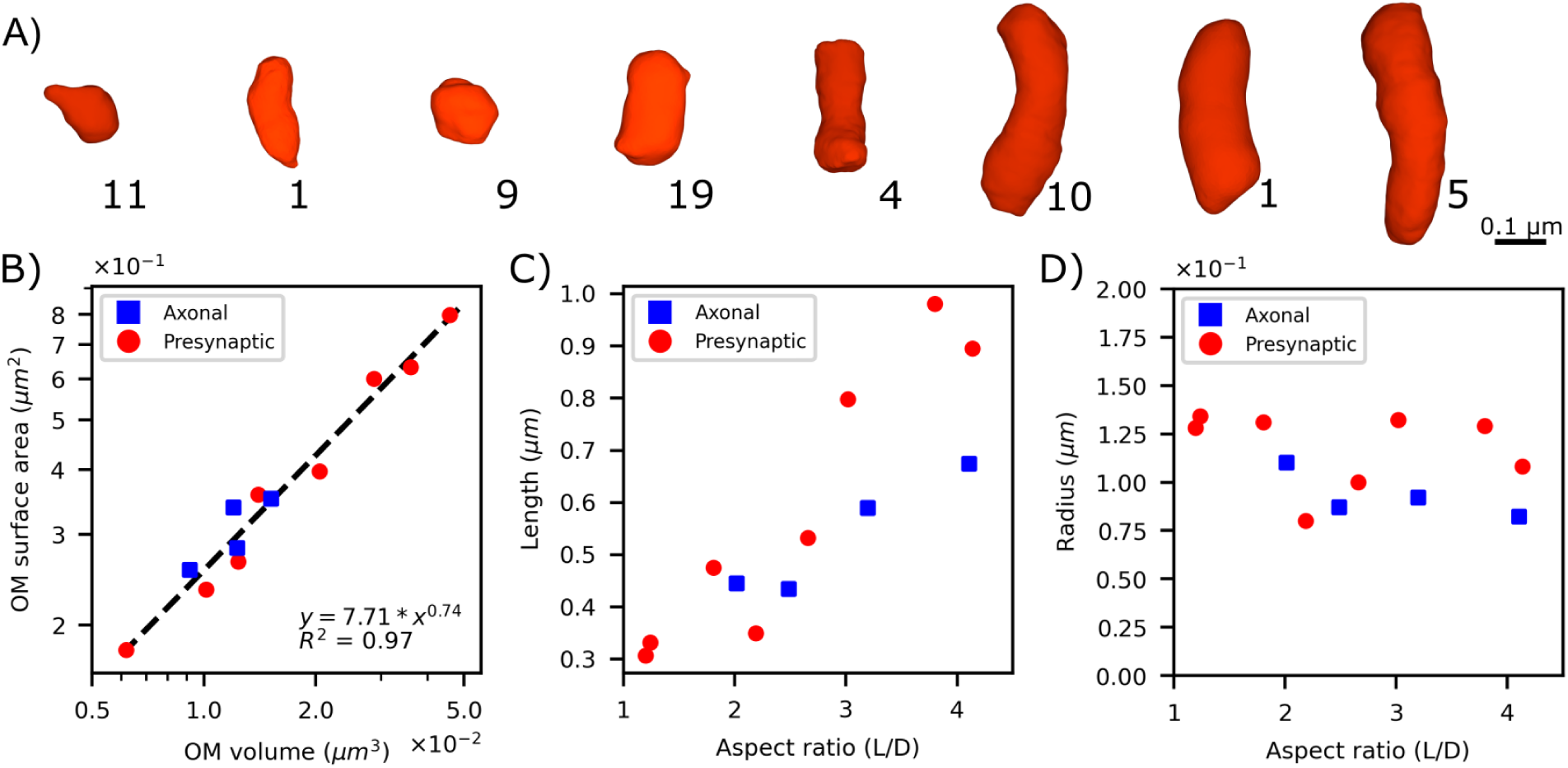
Wide range variation of reconstructed mitochondrial volumes, and their correlation with spatial location. Eight representative reconstructions of the outer membrane. Beside each shape a reference number is given to label each organelle. (B) OM membrane surface area correlates with the volume enclosed by the OM, axonal mitochondria (in blue) and presynaptic mitochondria (in red). (C) The length of the organelles increases with the aspect ratio. D) The mean radius of the organelles presents some variability for larger aspect ratio.

The shape of the outer membrane also varies within our data set, from globular to significantly elongated (Figure 2A). To quantify this observation, we calculated the aspect ratio of the shape. We defined the aspect ratio as the length of the reconstructed mesh divided by its width, such that elongated shapes are expected to have a higher aspect ratio and globular shapes are expected to have aspect ratio values close to one (Figure 2C). The width used to compute the aspect ratio are averaged quantities measured along the length of a mitochondrion, using a generated skeleton of the mesh (Methods). We contrasted the OM aspect ratio with the radius (Figure 2D) and found some variability on the organelles’ radius for longer organelles, with an average value of (0.12 ± 0.02) μm.

We grouped the mitochondrial reconstructions into two categories based on two geometric properties – the aspect ratio and the average principal curvature (refer to the next section for further details). The populations were identified using the K-means clustering algorithm. For each group, we calculated means and variances of five key geometric features – volume, surface area, radius, length and aspect ratio, see Table 1. The globular group has smaller aspect ratio (mean value 1.9) than the elongated one (mean value 3.44), with statistically different aspect ratio values (p-value < 0.026). As expected, the globular group is also shorter in length than the elongated one (although this difference is not significant). Some of the elongated organelles are also thinner, and for this reason the volume, albeit larger, is not significantly different than in the globular population (Table 1).

**Table 1:** Morphological properties of the outer membrane. Specific measurements were performed for each reconstruction. Averages and standard deviations over two distinct populations of mitochondria are presented. Two clusters were determined based on the aspect ratio and average of the first principal curvature of the OM (see methods for details).

| Property | Globular (N=6) | Elongated (N=6) | p-value |
| --- | --- | --- | --- |
| Aspect ratio | $1.9 \pm 0.6$ | $3.4 \pm 0.7$ | 0.026** |
| Length ($\mu\text{m}$) | $0.5 \pm 0.2$ | $0.7 \pm 0.2$ | 0.143 |
| Width ($\mu\text{m}$) | $0.24 \pm 0.04$ | $0.20 \pm 0.03$ | 0.143 |
| Volume ($\mu\text{m}^3$) | $0.02 \pm 0.01$ | $0.02 \pm 0.01$ | 0.931 |
| Surface area ($\mu\text{m}^2$) | $0.3 \pm 0.2$ | $0.5 \pm 0.2$ | 0.474 |
| Average first principal curvature (k1) ($\mu\text{m}^{-1}$) | $-0.3 \pm 0.5$ | $3 \pm 1$ | 0.002** |
| Average second principal curvature (k2) ($\mu\text{m}^{-1}$) | $-14 \pm 3$ | $-16 \pm 3$ | 0.474 |

\* $p < 0.05$ , \*\* $p < 0.01$

### Outer membrane curvature

To fully characterize the structure of mitochondria we employed tools from differential geometry [49, 22]. To identify their metabolic states, we quantified the curvatures of the surface reconstructions of the mitochondrial membranes.

For a given membrane, the principal curvatures characterize the major geometric properties and allow for the analyses of different structural motifs. The first principal curvature (k1) represents the maximum curvature at a given point on a surface and the second principal curvature (k2) corresponds to the minimum curvature [49, 22]. We use the principal components and their independent combinations; the mean curvature, given by *H* = (*k*1 + *k*2)/2, and the Gaussian curvature, by *K* = *k*1 × *k*2, to perform curvature analysis of the reconstructed membranes. The analysis was carried out with GAMer 2 [23, 24], using the Meyer, Desbrun, Schröder, and Barr (MDSB) algorithm [30] for curvature calculations. Positive curvatures occur when the outward facing normal and the curvature have the same directions, i.e. at concave regions.

To quantitatively understand what the curvature distributions of the OM mean, we compared them to the curvature distributions of idealized geometries. We consider spherical and capsule-like shapes (consisting of a cylinder with hemispherical ends), resembling approximations of globular and elongated geometries respectively. Both principal curvatures of a sphere of radius R are equal to 1/R, and negative with the sign convention applied (Figure 3A-B). For the capsule-like shape the principal curvature has contributions from the sphere (1/R) and zero from the cylinder (Figure 3A-B) while the second principal curvature receives contributions 1/R from all the vertices.

**Figure 3:**
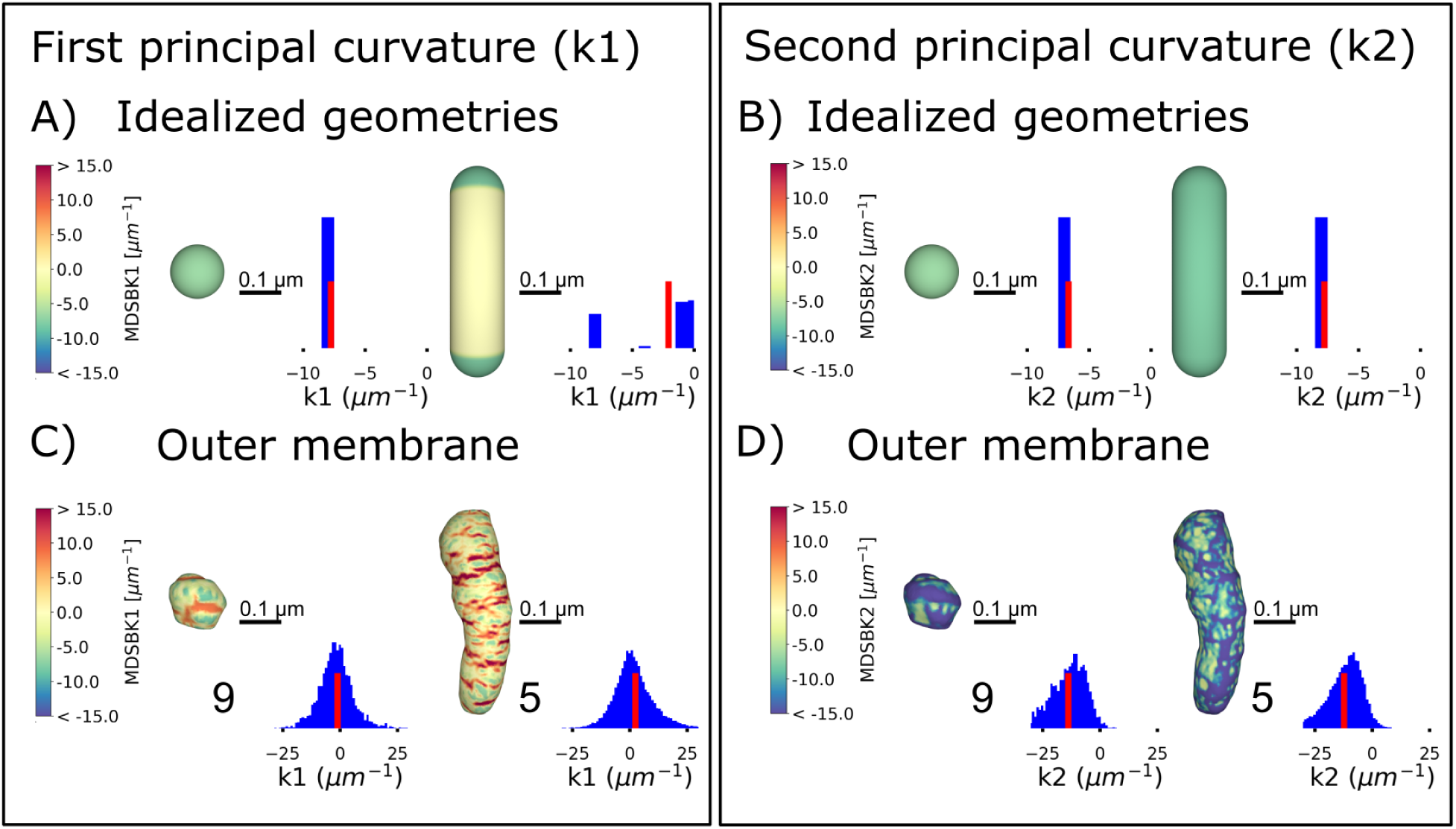
Curvature analysis of idealized geometries and the outer membrane. Heat maps of the first principal curvature k1, representing the maximum curvature at each vertex in the mesh for (A) a spherical shape and a capsule-like shape (composed of a cylinder and two hemispheres), and their respective curvature distributions. The sphere has a radius of 0.13 μm, thus its curvature is approximately −7.7 μm^−1^. An histogram of the curvature values is represented beside the shapes, with a red bar representing the average curvature of the distribution. The radius of the cylinder is 0.13 μm and its length is 0.98 μm. Therefore, the first curvature is approximately −7.7 μm^−1^ at vertices belonging to the semi-spheres and zero at the cylinder. (B) For the same shapes, heat maps of the second principal curvature k2, corresponding to the minimum curvature at each vertex in the mesh. For a sphere the second curvature is equal to the first, 1/R, −7.7 μm^−1^ in this example. For the capsule-like shape the contribution of all vertices is similar and equal to 1/R, −7.7 μm^−1^. (C) Heat maps of the first principal curvatures for two reconstructed outer membranes, representatives of the globular and elongated populations respectively. The curvature values mapped are between the 5th and 95th percentiles. The distribution of k1 values is positive and negative, showing different membrane concavities, positive values correspond with concave regions and negative ones with convex regions. An histogram of the curvature values is represented beside the shapes, with a red bar representing the average curvature of the distribution. D) Heat map of the second principal curvature for the same outer membrane reconstructions. Values represented are between the 5th and 95th percentiles. The distribution of k2 values is mostly negative. All curvatures were calculated with the MDSB algorithm implemented in GAMer 2. Color bars represent values of the curvatures in units of μm^−1^. Scale bar 0.1 μm.

For the idealized geometries considered here, the average first principal curvature is smaller for the sphere and larger for the capsule-like shape, whereas the second principal curvature is equal for both geometries (the average of the distributions is represented with a red bar in Figure 3). The average first principal curvature in the capsule-like shape is greater than the sphere due to the contributions of the flat cylindrical regions (where *k*_1_ = 0) to the average. Using the insight provided by the average principal curvatures, we can thus differentiate between globular and elongated geometries.

In Figure 3C-D we show two representative reconstructions of the outer membrane, with their respective first and second curvature distributions mapped to the membrane. The first curvature takes positive and negative values presenting areas of different concavity, dominantly the vertices with negative curvature occur at the ends of the shape, at convex regions, while the positive ones are distributed at the central part. Similarly to the idealized geometries, the average first principal curvature is smaller for the globular shape (Figure 3C-D). On the contrary, the average second principal curvatures are similar.

We clustered the reconstructions based on the values of the aspect ratio and the average first principal curvature using the K-means algorithm (Figure 4A). The globular population has an average k1 and aspect ratio significantly smaller than the elongated population (p-value < 0.004 and < 0.026 respectively), while the average k2 is not significantly different between both groups, as expected from the analysis with the idealized shapes (Table 1 and Figure 4B). The globular set is mostly composed of presynaptic mitochondria, 5 organelles out of 6 of the organelles are located at boutons. On the other hand, the elongated group is composed of a mixture of axonal (N = 3) and presynaptic mitochondria (N = 3).

**Figure 4:**
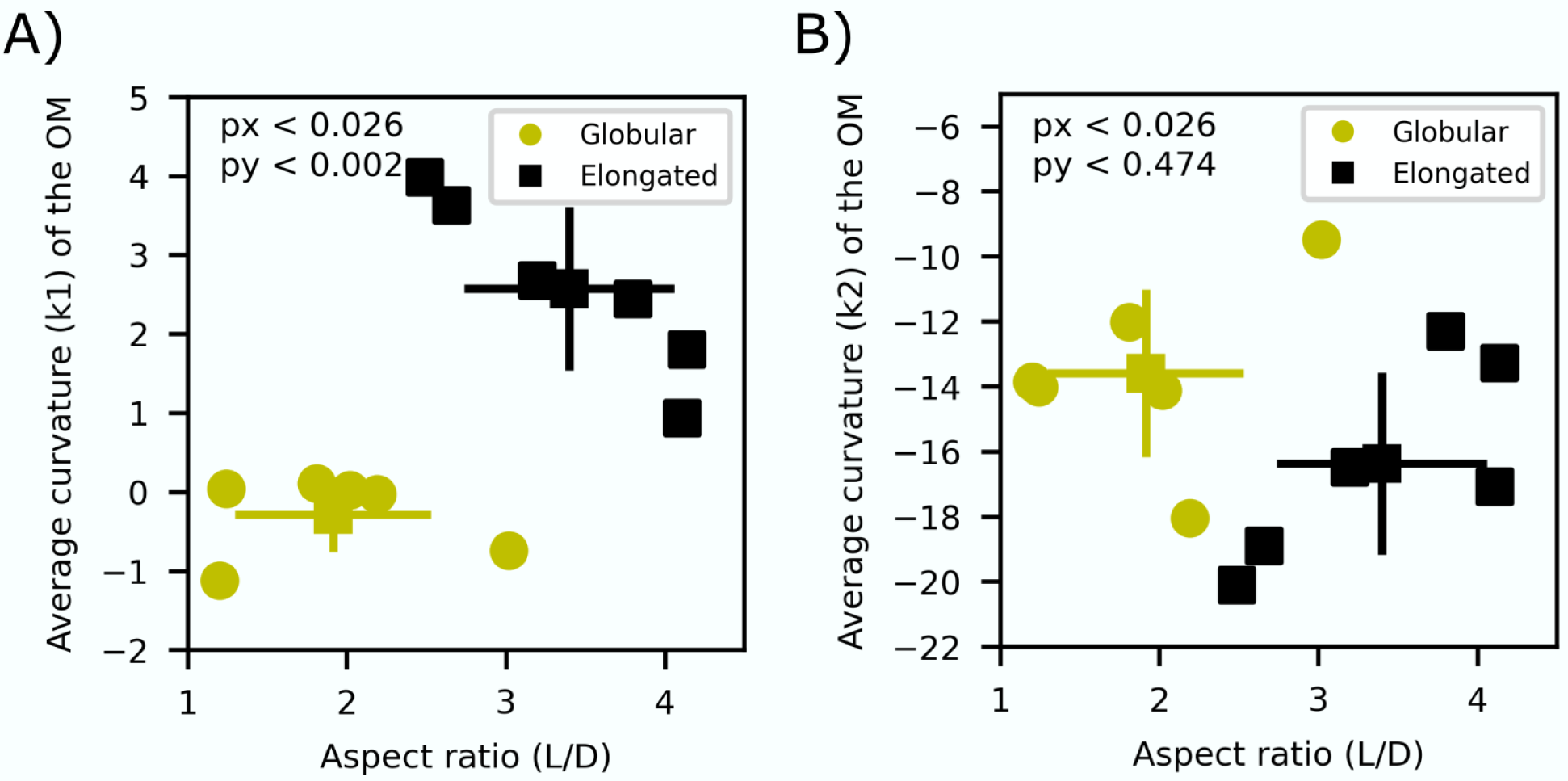
Two populations of organelles are identified using the principal curvature k1 and the aspect ratio as key features. (A) Average principal curvature (k1) of the outer membrane and aspect ratio of the shape. Organelles are separated into two clusters employing the K-means clustering algorithm. The globular population (yellow circles) has a smaller aspect ratio and average first principal curvature while the elongated one (black squares) has greater aspect ratio and average first curvature. The mean values and standard deviations are represented for each population. Both populations are statistically significantly different in regards to both variables (with p-values < 0.01 in both axis). (B) Average second principal curvature and aspect ratio for both identified populations. The second principal curvature is similar for both populations (p-value < 0.6) as expected comparing with the idealized shapes (refer to main text for further details).

Interestingly, a small set of the organelles (N=4) present protrusions in the OM. In all cases, an area of positive and high curvature at the neck of the protrusion is followed by an area of high negative curvature (Supplementary Figure 10). These organelles have an average radius of (0.17 ± 0.01) μm which is significantly smaller than for the rest of the group (0.24 ± 0.02) μm, p-value < 0.004). The average length of these two groups is similar, and the volume is smaller for the ones with protrusion with (0.011 ± 0.003) μm^3^ compare to (0.02 ± 0.01) μm^3^ respectively, although this difference is not significant (p-value < 0.2). These measurements not only give us quantitative insights into the geometric features of the OM but, as we will discuss later, can also inform the design of geometries in spatial simulations.

### Geometric features of the inner membrane

We next analyzed the geometric features of the inner mitochondrial membrane. The ultrastructure of neuronal mitochondria is composed of a complex network of lamellar and tubular cristae (Figure 5A-B), which generates a number of sub-compartments as the intracristal space (ICS), intermembrane space (IMS) space and the matrix (Figure 1A). The ICS is encapsulated by the cristae membrane, opening into the inter membrane space (between the IBM and OM) via CJs. Figure 5A-B shows five reconstructions of the cristae membrane along with three of their skeletons. We first measured global features of the inner membrane, and its sub-compartments as the surface areas and volumes, and then quantified the number of cristae junctions on the membrane for each of the nine reconstructed mitochondria. Based on these measurements, we determined various ratios of surface areas and volumes of the mitochondria ultrastructure to facilitate comparison with the literature (Table 2). We assessed features of the membrane such as its continuity, and analyzed the motif composition using tools from differential geometry.

**Figure 5:**
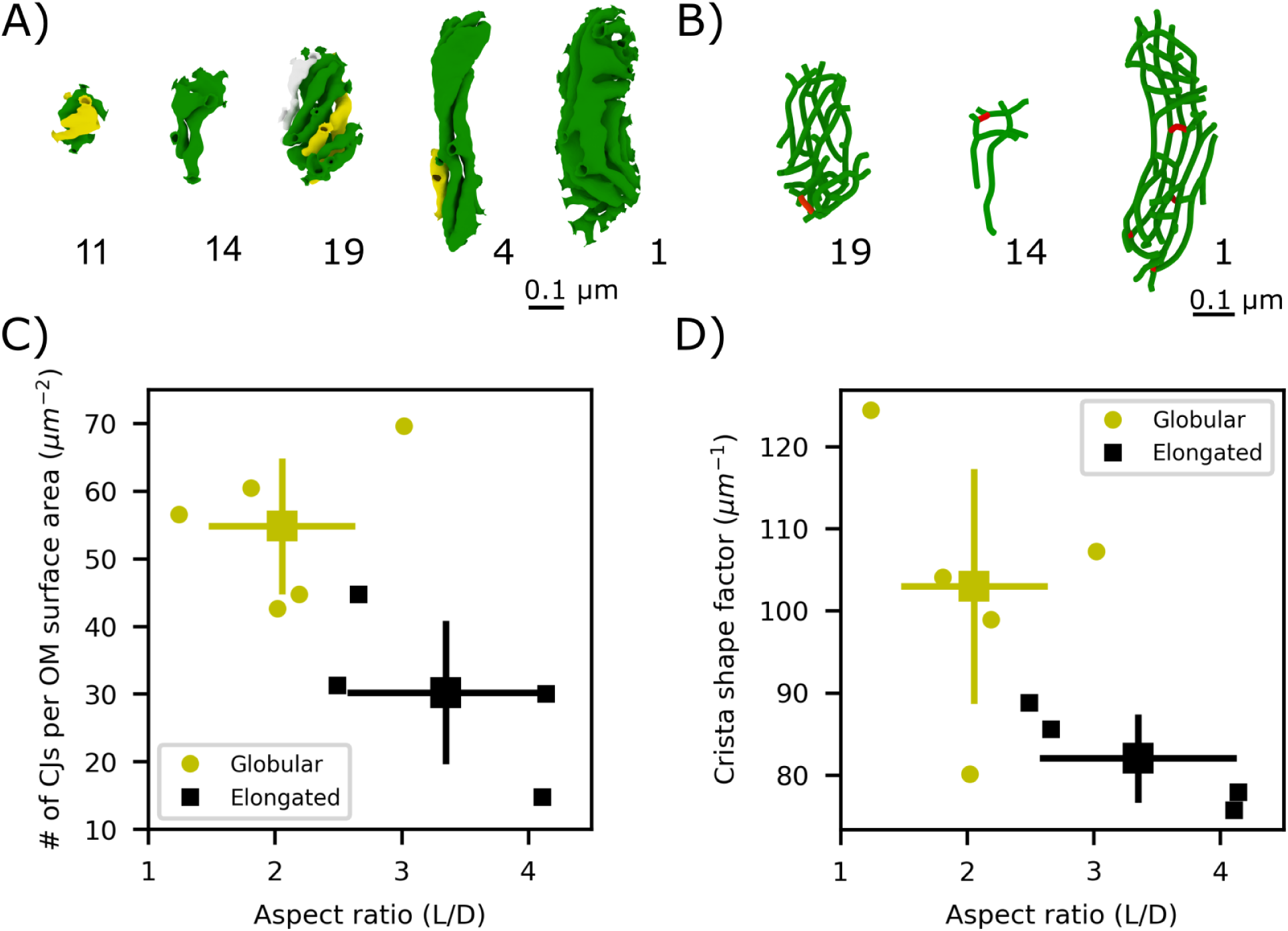
Morphological characterization of the internal structure for nine mitochondrial inner membrane reconstructions. The cristae membranes form a complex network of tubular and laminar structures. (A) Five cristae membrane reconstructions from the mitochondria indicated. A different color is used for each separate, non-interconnected cristae section (B) Three skeletons of the cristae membrane from the indicated mitochondria (see Methods). Red indicates connected segments between compartments. (C) The density of cristae junctions, defined as the # of CJs per OM surface area, is greater for globular organelles (yellow circles, p-value < 0.079). The mean values and standard deviations bars of are represented for each population. (D) The cristae surface area-to-volume ratio, also known as the crista shape factor, is also larger for globular organelles (p-value < 0.079). The mean values and standard deviations bars of each cluster are represented.

**Table 2:**
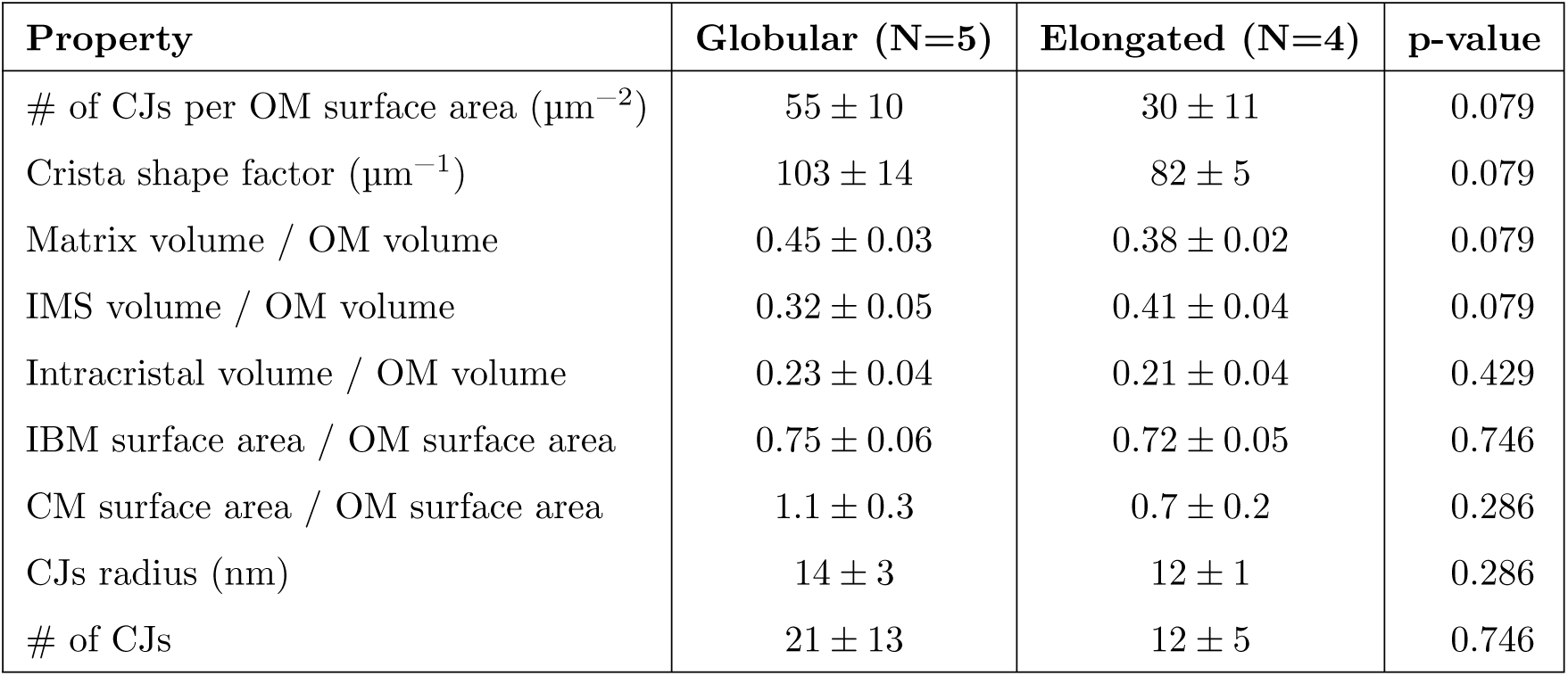
Morphological properties of the inner membrane for both populations identified with the aspect ratio and mean curvature of the outer membrane (Table 1); averages and standard deviations over each population are presented.

The variability of the density of cristae junctions and of the cristae membrane surface area-to-volume ratio, also called crista shape factor in the literature [37], with the aspect ratio was analyzed (Figure 5C-D). We found that both quantities are larger for globular organelles, i.e. organelles with smaller aspect ratio (Figure 5C-D). The larger density of CJs for globular organelles can be explained by the smaller OM surface area of globular organelles and the larger number of cristae junctions found in globular organelles (Table 2). Moreover, the spatial distribution of the junctions is not homogeneous, and seems to concentrate at the ends of the elongated sampled mitochondria. The density of the CJs found here (55 and 30 per μm^−1^ for globular and elongated populations respectively) are smaller than the values previously reported in the literature for presynaptic cerebellar mitochondria (136 μm^−2^) [40].

Given the high quality of the reconstructed meshes, we also quantified the surface areas and volumes of the different organelle’s sub-compartments. We calculated the total volume taken up by each of the three compartments: the intracristal space, the intermembrane space and the matrix, relative to the OM volume. Averaged quantities are presented in Table 2, for the globular and elongated populations previously identified. Our measurements are consistent with previous work reporting crista to outer membrane surface area ratio of ~1 for presynaptic cerebellar mitochondria [40].

Multiple tubular cristae near the cristae junctions are seen connecting to form larger, lamellar compartments in the center of the inner membrane, consistent with other neuronal mitochondria reconstructions [39]. We determined different aspects of these cristae networks by analyzing their continuity. All cristae in the mitochondria reconstructed here are connected to the inner boundary membrane through cristae junctions, meaning that there are no self-contained compartments of intracristal space. Therefore, we classified the cristae membrane which forms these compartments as contiguous with the inner boundary membrane. We further explored the inter-connectivity within the cristae membrane (disregarding now the inner boundary membrane). These can be described as different cases where the cristae membrane is dis-contiguous with itself. In one third (three) of the nine inner membrane reconstructions, the entire intracristal space forms a single, fully connected compartment, and so does the cristae membrane. In another third there are three disconnected intracristal spaces, and in the remaining third of the samples there are three disconnected intracristal spaces. In Figure 5A, we colored the cristae with the same color when the cristae membrane is contiguous (as in reconstruction # 14 and # 1 in Figure 5A), and we used a different color to signal when the membrane is discontinuous. The continuity of the cristae were based on our analysis of the reconstructed meshes using tools in Blender (refer to Methods section). However, to further identify the connections between different cristae, we generated skeletons of the cristae membrane (Figure 5B), and highlighted in red the tubular structures connecting different cristae compartments. This level of connectivity between membranes and the formed compartments could have significant implications for molecular diffusion as we discuss in the next section.

In 5 out of the 9 cristae membrane reconstructions, we also observed that lamellar cristae sheets form corkscrew-like twists which were always right-handed. Two examples are reconstructions # 19 and # 4 in Figure 5A. These structures are similar to the distribution of cristae junctions that were recently shown in yeast and cultured mammalian cell lines [47]. Also, the spatial distribution of cristae junctions is irregular, and seems to concentrate at the ends of the elongated mitochondria, as already reported in the literature [39]. Cristae structures with a helicoidal twist appear to have a higher cristae junction density on the top and bottom of the twist compared to the center.

Another important geometric property of an orientable surface is its genus; the genus of an orientable surface is the number of holes it has. As an example, a sphere has genus zero, while a torus has a genus of 1. We evaluated this property for all the reconstructed inner membranes, and found a large variability, with values ranging from 4 to 47 (refer to Methods Section for details). The largest value is found in the reconstruction with the most cristae junctions and connected inner membrane (reconstruction # 1 in Figure 5A), while the smallest value is found in the organelle with the least cristae junctions (reconstruction # 7), these two properties are highly correlated Supplementary Figure 12. Interestingly, all the organelles with protrusions (n = 3) have an average genus of 5, whereas those without protrusions have an average genus of 22 (p-value < 0.02).

### Inner membrane curvature

To dissect the structural components of the complex network of cristae, we performed curvature analysis of the inner mitochondrial membrane. Given the intricate architecture of the inner membrane, in addition to quantifying the curvature, we also developed tools to quantify geometric motifs on the membrane.

To begin, we calculated the first principal curvature at each vertex of the inner membrane and color coded its values in a heat map representation (Figure 6A), for values between the 5th and 95th percentiles. For visualization purposes, only the cristae membrane is shown; however, the entire inner membrane is analyzed to preserve the curvature at each vertex. The distribution of the first curvature is primarily positive, with the highest values found at cristae junctions, at the connection site between cristae, branch sites of cristae, at the edge of lamellar cristae and along the length in tubular cristae. Similarly, we generated a heat map of the second principal curvature of the inner membrane, representing the minimum curvature at each vertex (Figure 6B), for values between the 5th and 95th percentiles. The values of k2 are mainly negative, and particularly high at cristae junctions, connections between cristae, and at the branch sites of cristae, complementing in some regions the high and positive k1 curvature. Remarkably, there are also small areas of positive curvature at the ridge of cristae, additionally quantified as geometric motifs.

**Figure 6:**
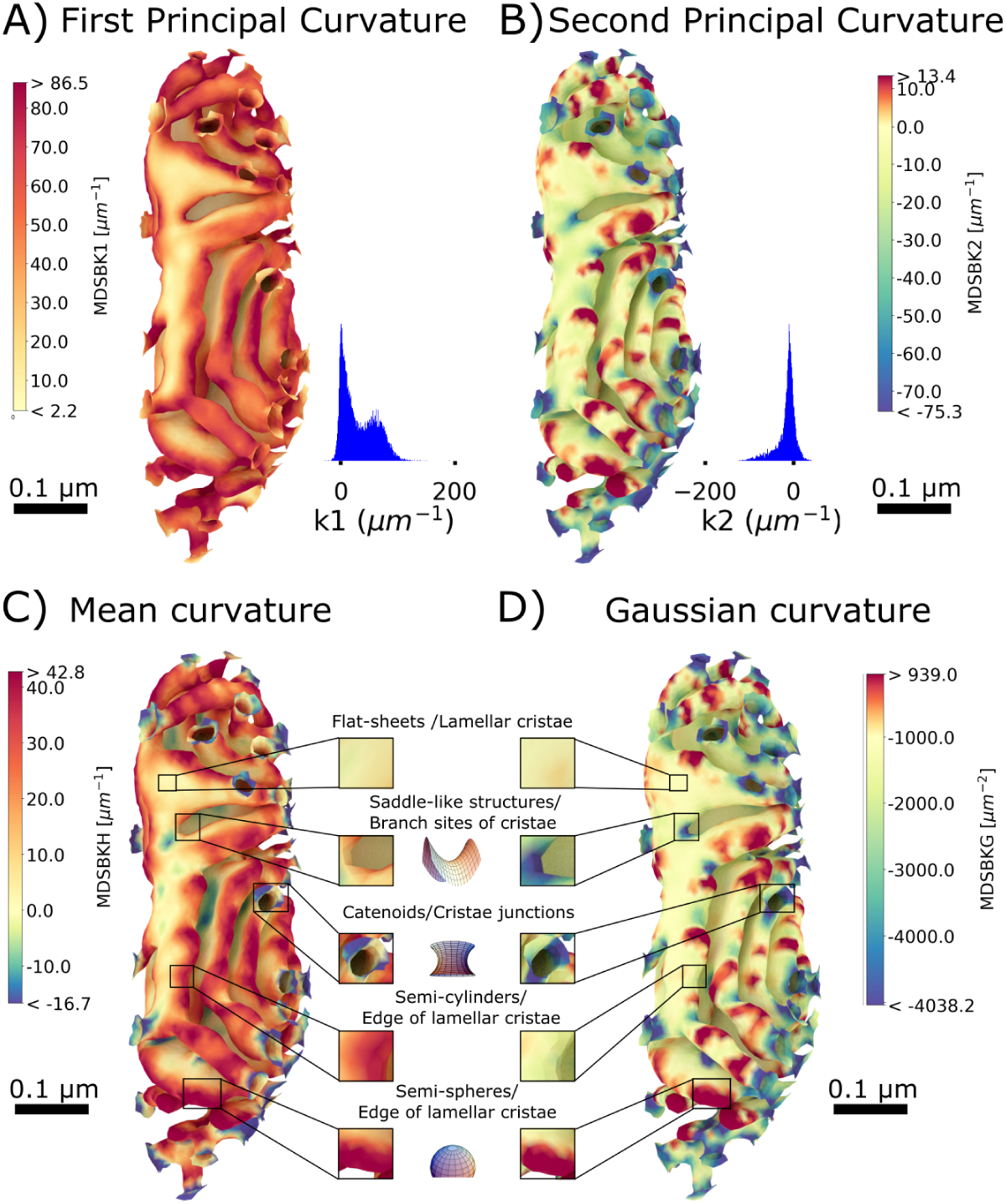
Curvature analysis of the inner membrane of reconstruction # 1. (A) Curvature heat map of the first principal curvature k1, representing the maximum curvature at each vertex in the mesh (only the cristae membrane is represented), for values between the 5th and 95th percentiles, and their respective distribution values. The distribution of k1 is mostly positive, with the highest values found at cristae junctions, the connection site between cristae, branch sites of cristae, at the edge of lamellar cristae and along the length in tubular cristae. B) Curvature heat map of the second principal curvature k2, corresponding to the minimum curvature at each vertex in the mesh, for values between the 5th and 95th percentiles. The distribution of k2 is mostly negative, and particularly high at cristae junctions, connections between cristae, and at the branch sites of cristae, complementing the high positive curvature k1 as expected for minimal surfaces. (C) Heat map of the mean curvature, corresponding to the average value of the first k1 and second curvature k2, for values between the 5th and 95th percentiles. (D) Heat map of the Gaussian curvature, corresponding to the product of first and second curvature, for values between the 5th and 95th percentiles. Four characteristic structural motifs emerged in the cristae membrane: flat-sheets are in lamellar cristae with mean and Gaussian curvatures approximately zero; saddle-like structures with zero mean and nonzero Gaussian curvature are at cristae junctions and branch sites of cristae; at the edge of lamellar cristae we found two characteristic shapes: semi-spherical like shapes with nonzero mean and Gaussian curvature and semi-cylindrical shapes with nonzero mean and zero Gaussian curvature. Curvatures were calculated with the MDSB algorithm implemented in GAMer 2. Color bars represent values of the curvatures in units of μm^−2^ for the Gaussian curvature and μm^−1^ for all the rest, Scale bars 0.1 μm. Structural motif representations were generated with different notebooks from MathWorld [56, 53, 55, 54].

### Structural motifs in the inner mitochondrial membrane

To further assess the presence of structural motifs on the cristae membrane, we calculated the mean curvature H and the Gaussian curvatures K at each vertex of the inner membrane reconstructions (Figure 6C-D), corresponding to the mean and product of the principal curvatures, respectively. Across all the cristae membranes analyzed, we found four characteristic shapes in the cristae membrane, corresponding to different combinations of the mean (H) and Gaussian curvature (K) [49, 22]: flat-sheets for which both curvatures vanish (H = 0 and K = 0), saddle-like structures (H = 0 and K ≠ 0), semi-spherical shapes (H > 0 and K ≠ 0) and semi-cylindrical shapes (H ≠ 0 and K = 0).

Flat-sheets are present in lamellar cristae, with mean and Gaussian curvature approximately zero (for all quantitative analysis we consider ~ 0 mean curvatures smaller in absolute value than 10 μm^−1^, and Gaussian curvatures smaller in absolute value than 300 μm^−2^). Surprisingly, we found a multitude of saddle-like structures with zero mean and nonzero Gaussian curvature at cristae junctions, branch sites of cristae, and in some areas of the lamella (Figure 6C-D). In particular, the structures found at cristae junctions are reminiscent of catenoids (with H = 0 and K < 0). These types of neck-like structures form connections between regions of different curvatures as was discussed in [5]. Flat-sheets and saddle-like structures are made entirely of saddle points where two opposite curvatures generate a mean curvature of zero. The presence of these structures are reminiscent of minimal surfaces in differential geometry and have been noted in the endoplasmic reticulum (ER) reconstructions as well [51, 28].

At the edges of lamellar and tubular cristae, we found two characteristic shapes: half-spherical shapes and half-cylindrical shapes. The spherical regions have high mean and Gaussian curvature values (Figure 6C-D). The Gaussian curvature, defined as the product of the two principal curvatures, is therefore positive when the principal curvatures have the same sign, as found in these regions. In contrast, the Gaussian curvature vanishes in cylindrical shapes, as found in some regions of lamellar and tubular cristae (Figure 6C-D).

Thus, using our curvature quantification, we have classified four structural motifs: flat-sheets, saddle-like structures, semi-cylindrical egdes of lamellar cristae, and hemispherical edges of lamellar cristae. We quantified the presence of the aforementioned four structural motifs in all the reconstructions by computing the fraction of the cristae membrane with mean and Gaussian curvatures similar to their respective idealized values, to the total cristae membrane surface area (see Table 3; representations of these surfaces for one reconstruction are shown in Supplementary Figure 13). Remarkably, the percentage of flat-sheets and spherical shapes is not significantly different between the globular and elongated populations - 12% and 11% respectively (Table 3). However, the presence of saddle-like structures consists of approximately 4% of the cristae membrane with a statistical significance of (p-value < 0.016) difference of 2.3% for the globular and 3.7% for the elongated populations. The presence of semi-cylindrical shapes is also different between the two populations, with larger portions of the membrane occupied by this structural motif in elongated mitochondria (Table 3).

**Table 3:** Quantification of the presence of four structural motifs for both globular and elongated mitochondria populations (see Table 1). Averages and standard deviations over each population are shown for the % cristae membrane surface area occupied by each motif. The last entry is the percentage of the cristae membrane with vertices whose first principal curvature k1 exceeds 80 μm^−1^.

| Property | Globular (N=5) | Elongated (N=4) | p-value |
| --- | --- | --- | --- |
| % flat-sheets | $12 \pm 10$ | $11 \pm 4$ | 0.746 |
| % saddle-like structures | $2.3 \pm 0.6$ | $3.7 \pm 0.2$ | 0.016* |
| % semi-cylindrical shapes | $4.64 \pm 0.02$ | $5.65 \pm 0.03$ | 0.079 |
| % semi-spherical shapes | $13 \pm 5$ | $13 \pm 2$ | 0.746 |
| % of the area with max k1 ($> 80 \mu\text{m}^{-1}$) | $13 \pm 6$ | $5 \pm 3$ | 0.079 |

We quantified the area of cristae membrane formed by vertices with intense curvature. To clearly visualize these regions, we generated heat maps of the first principal curvature color coding only the curvature values larger than 50 μm^−1^ for the globular and elongated populations (Figure 7A and B, respectively). Additionally, we analyzed the correlation between the fraction of the area formed by intense curvature regions and the aspect ratio, in both populations (Figure 7C). We found on average 13% of the cristae membrane is formed by vertices with high curvature in globular organelles, whereas in elongated mitochondria this area represents 5% of the total cristae membrane (Table 3). Since ATP synthases accumulate at regions of most intense curvature, we consider this area as a proxy of the energetic capabilities of a given mitochondrion. Therefore, our results suggest that globular organelles show features of high energy capacity.

**Figure 7:**
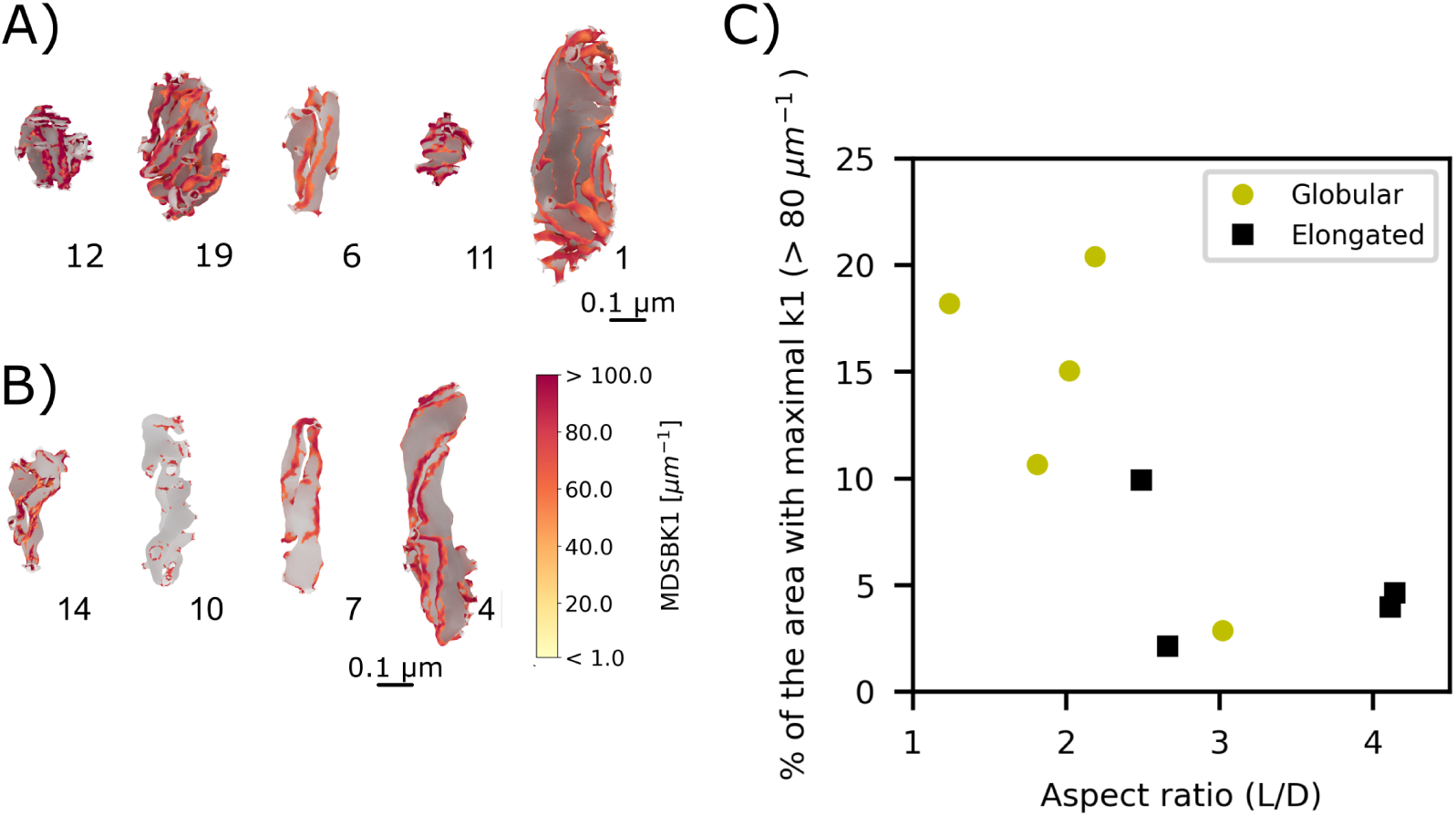
High curvature regions of the inner mitochondrial membrane. A-B) Only vertices of the cristae membrane with the first principal curvature k1 greater than 50 μm^−1^ are shown in color according to the color bar. Remaining vertices are in grey. (A) Globular mitochondria population and (B) Elongated mitochondria population. C) Percentage of the surface area formed by vertices of the cristae membrane with first principal curvature k1 greater than 80 μm^−1^, as a function of the outer membrane aspect ratio for the globular (yellow circles) and the elongated (black squares) populations.

## Discussion

Brain activity is accompanied by a large energy demand and therefore relies on efficient energy metabolism. In particular, neurons and their synaptic connections are energetic ‘hot spots’ and predominantly rely on mitochondrial ATP production. For this purpose, presynaptic mitochondria seem to exhibit a specialized morphology but their functional consequence is still not fully understood. Here, we used electron tomography to investigate structural properties of these specialized organelles. Electron tomography has proven to be a powerful tool to obtain high resolution images of *in situ* organelles [13] and particularly useful to resolve the presynaptic structure [36]. The final resolution of electron tomograms depends on several factors, including sample preparation, section thickness and electron microscope voltage. In this work, samples were high-pressure frozen followed by freeze-substitution with fixatives (HPF-FS) [45], producing a large volume of a well-preserved tissue that is subsequently sectioned with minimal material loss. This combination of factors generates serial tomograms with an isotropic voxel size of 1.6 nm, that are employed to segment and reconstruct mitochondria from axons and presynaptic terminals. High quality whole organelle reconstructions produced with the developed workflow, that includes new mesh processing software GAMer 2 [23, 24] and new alignment methods [57], not only allow for nuanced geometric characterization of the internal architecture, as presented in this work, but also provides *in silico* representations for spatial computational modeling [14]. This type of reconstructions and quantifications are hard to conduct on a routine basis because of the technical challenges involved. Here, we have overcome these challenges with newly developed methods. Such interdisciplinary effort lays the foundation for future studies that can map internal mitochondrial physiology, energy production, and metabolism in neurons.

Our analysis reveals that the 12 mitochondria we analyzed can be categorized as elongated or globular. Pro-found architectural differences between globular and elongated mitochondria, are not only reflected in the external structure but also in the internal architecture (Figure 5). We found that globular organelles have higher density of cristae junctions and surface area-to-volume ratio, and notably the percentage of the area formed by vertices with extreme curvature is larger in these organelles (Figure 7C). Since dimeric complexes of ATP synthases accumulate in regions with high curvature of the cristae membrane and form ribbon structures which enforce that curvature [48, 33], the increment in high curvature regions in our globular reconstructions suggests that globular organelles, show features of high energy capacity.

Some of the features reported here have been measured previously for axonal and presynaptic mitochondria like the lengths, widths, cristae junction diameters [35, 27, 43], crista shape factors [37, 7], density of cristae junctions [39, 40] and are in reasonable agreement with the values reported in the literature. In particular, a high crista shape factor have been already associated with organelles located at high energy demanding locations [37, 7]. In addition to the above listed features, we also calculated the principal curvatures for the membrane surfaces and found important structural motifs. Such calculations have implications in our understanding of disease states. For example, near total loss of ATP synthase dimerization due to a neurodegenerative mutation causes profound disturbances of mitochondrial crista ultrastructure in fibroblast cultures derived from a Leigh syndrome patient [43]. One of the key architectural features different than in control cells is an increment in the angle of curvature at the tips of the cristae. This feature relates to our curvature measurement of the inner membrane. Another feature of the patient mitochondria is reduced crista surface area-to-volume ratio [43]. Further investigation with the here developed methods would allow for a deeper understanding of potentially impaired mechanisms.

The ultrastructure of neuronal mitochondria forms a complex geometric structure. We evaluated features of this membrane which were inaccessible before, such as its continuity. We used skeletons of the reconstructions to automatically measure structural features and also to identify the connections between different cristae. Contiguous compartments can share resources, molecules and proteins. They can potentially reach all adjacent spaces diffusing through the connected volumes or membranes. These possibilities and their functional consequences can be further explored with computational modeling by quantifying the diffusion capacity within the membranes and in the confined 3D space.

New tools to estimate the curvature at each vertex in the reconstructions (implemented in GAMer 2) allowed us to quantify features of the membranes that would remain as qualitative observations otherwise. We used the curvature to objectively separate the organelles into two distinct sets, to identify protrusions in the membrane, and to quantify the presence of structural motifs in the cristae membrane. Beside the large variability found in the cristae membranes, the percentage of flat-sheets and spherical motifs is on average the same in both populations. However, this is not the case for saddle-like structures and semi-cylinders.

The structural motifs identified in our analysis of membrane surfaces suggest that the geometry of mitochondrial membranes may be understood from mechanical reasoning. The shape of a cellular membrane can be thought to be a minimum energy state for the bending energy of the lipid bilayer [18]. Several mechanisms can generate positive or negative curvature in membranes [29] including changes in the lipid composition, cytoskeletal proteins, scaffolding by external membrane proteins and influence of integral membrane proteins. Semi-cylindrical and semi-spherical shapes, as found in mitochondria, would require high membrane asymmetry of lipid composition to be maintained [58]. It is highly likely that the crowded mitochondrial milieu, combination of peripheral and integral membrane proteins coupled with external forces play a role in generating these membrane shapes.

One curious feature about our observations is the large fraction of flat sheets and saddle-like structures, which are minimal surfaces, so-called because they minimize a surface energy associated with curvature. It is at its lowest at saddle points, where two opposing curves of equal intensity form a saddle-like shape [1, 5].

The classic illustration of minimal surfaces shows a soap film stretched along a wire frame. These soap films, much like cellular membranes, are lipid structures which will take whatever shape that will minimize the surface energy. This propensity for membranes to form minimal surfaces has been demonstrated using the catenoid shape of neck-like structures which form during fission and fusion as well as in endo- and exocytosis [5]. As seen with the wire frame required to create a minimal surface from a soap film, membranes require structural elements around which these low-energy shapes can develop. A number of proteins have been identified as key regulators of cristae organization and remodelling, among them OPA1, MICOS complex and ATP synthase have been recognized as the main regulators [15]. And in particular OPA1 and MICOs complexes have been shown to regulate cristae junctions [46, 20]. These protein complexes can potentially act as the scaffolds or integral proteins applying the forces required to bend the membrane.

To summarize, we have quantified multiple morphological features of presynaptic mitochondria using end-to-end reconstructions. Analysis of these mitochondria have revealed common principles of organization across the sampled population suggesting that there exist design criteria for healthy mitochondria.

Recently, a series of studies on synaptic geometry and structural plasticity have been conducted by us and others [4, 3, 26, 8]. Parallel studies have focused on the role of the ER and mitochondria in regulating the spine calcium and how internal organelles can alter the calcium and ATP dynamics [14, 25, 44]. We anticipate that the mitochondrial geometry features we describe here will now serve as a platform for future high-dimensional simulations of mitochondrial dynamics and quantitative mapping of the mitochondrial structure-neuronal energy landscape.

## Methods

Two serial tomographic data sets from the cerebellar neuropil of two mice yielded 8 presynaptic and 4 axonal mitochondria which were fully contained in the samples with a tomographically reconstructed isotropic voxel size of 1.64 nm. With physical serial sections of 300 nm each, mitochondria are contained in 1 to 3 physical sections. At the area between these sections, missing material of approximately 15 nm – 20 nm and distortion caused by the slicing process was corrected for using state-of-the-art image alignment [57] and gap interpolation techniques described later in this section. This allowed for the complete reconstruction of 12 of these mitochondria, for 9 of them we generated complete reconstructions of all the membranes.

High image resolution enables the contours of the OM, IBM, and cristae membrane of the mitochondria to be observed and manually traced on each virtual section (Figure 1C) using reconstruction software. The process of creating a three-dimensional surface mesh from the traced contours (Figure 1B) and smoothing out surface noise using mesh conditioning algorithms, resulted in reconstructed surfaces with less than 2 nm error in surface placement.

### Specimen Preparation

A mouse (C57BL/6NHsd, male, 1-month old) was anesthetized with ketamine / xylazine and fixed by transcardial perfusion. The vasculature was briefly flushed with oxygenated Ringer’s solution, followed by 2.5% glutaraldehyde, 2% formaldehyde, 2 mM CaCl_2_ in 0.15 M sodium cacodylate buffer for 10 minutes. The fixative solution was initially 37 °C and was cooled on ice during perfusion. The brain was removed from the cranium and placed in the same fixative solution for 1 hour at 4 °C. The cerebellar vermis was cut into 100 μm-thick sagittal slices on a vibrating microtome in ice-cold 0.15 M cacodylate buffer containing 2 mM CaCl_2_ and briefly stored in same buffer prior to HPF. A 1.2 mm tissue punch was used to take tissue from vibratome slices. Punches were placed into 100 μm-deep membrane carriers and tissue was surrounded with 20% bovine serum albumin (Sigma) in cacodylate buffer to prevent air pockets. The carrier was then frozen with a Leica EM PACT2. Freeze substitution was performed in a Leica AFS2 using extra dry acetone (Acros) as follows: 0.1% tannic acid at −90 °C for 24 hours, wash 3 × 20 min in acetone, 2% OsO_4_ / 0.1% uranyl acetate at −90 °C for 48 hours, warmed for 15 hours to −60 °C, held at −60 °C for 10 hours, and warmed to 0 °C over 16 hours. The specimen was infiltrated with a series of Durcupan ACM:acetone solutions at room temperature and then embedded in 100% Durcupan at 60 °C for 48 hours.

### Electron Tomography

A diamond knife was used to collect ribbons of 300 nm-thick serial sections on 50 nm-thick Luxel slot grids. Sections were glow discharged and coated by dipping grid briefly in 10 nm colloidal gold solution (Ted Pella). Serial tomographic reconstructions were generated of parallel fiber - Purkinje cell synapses using an FEI Titan 300 kV TEM equipped with a 4k x 4k CCD camera (Gatan Ultrascan). For each ROI, tilt series were collected with 0°, 45°,90°, and 135° degrees of in-plane specimen rotation. Each of these tilt series were collected from −60° to 60° with 1° tilt increments. Images were collected with a pixel size of 0.4 nm. Images were binned by 4 prior to tomographic reconstruction with TxBR [41]. If an ROI spanned multiple serial sections, a serial tomographic reconstruction was created.

### Alignment of Serial Tomographic Volumes

The serial tomographic datasets consist of 4 and 7 physical serial sections. Within each serial dataset, the tomographic reconstruction of each 300 nm-thick physical section results in a separate volume dataset for each physical section. We used image registration software, SWiFT-IR [57], to precisely align the reconstructions of the physical sections with one another. To accomplish this, alignment was performed between the last virtual section a given physical section and the first virtual section of the next physical section. No alignment was performed between the virtual sections within a physical section as the tomographic reconstruction produces virtual sections that are already aligned. It is important to note that the physical sections sustained a certain unknown amount of in-elastic deformation due to the cutting forces of the diamond knife during sectioning. Because of this deformation, we used SWiFT-IR to bring the sections into the best alignment possible using only 2D affine transformation implemented as X translation, Y translation, rotation, X scale, Y scale, and X skew. To further constrain the alignment transformation we noted that the determinant of the 2D affine transformation matrix can be interpreted as the scaling of the area of a unit square. After an initial alignment step we analyzed the determinant of the affine transform between each pair of physical sections and identified the physical section which required the least overall deformation of the area. We then defined this section as the alignment reference for the serial dataset by fixing the transform for this section to the identity transform and adjusting the remaining transforms in the dataset accordingly, yielding the final aligned volume spanning all the physical sections for each serial tomographic dataset.

### Manual Tracing of Mitochondria

The tomographic volumes were visualized in 3DEM software, Reconstruct [11] or IMOD [21], and the membrane-bounded structures of mitochondria were identified. The contours of the outer leaflet of the outer membrane, the inner leaflet of the inner boundary and cristae membrane were manually traced as separate objects. Although the inner membrane boundary and the cristae membrane form one contiguous structure in mitochondria, they were traced as two separate objects (Figure 1C) and later combined (Figure 1F). This procedure averts the meshing challenges posed by a single set of contours with deep invaginations. At cristae junctions, the inner boundary membrane object was traced as passing over the junction and the separate cristae membrane object was traced at the junction in a way that intersected with the inner boundary membrane while carefully following the curvature at the mouth of the junction before smoothly intersecting and protruding beyond the tracing of the inner boundary membrane. These protrusions acted as markers of the junctions to distinguish them from artifact and preserve the shape during the mesh smoothing process performed at later stages. Contours were traced of most clearly visible membrane leaflet for each object (the cytosol side of the OM and the matrix side of the IBM and cristae), with the other leaflet being added later. Three of mitochondria were traced multiple times to estimate the error due to manual tracing. We estimate the tracing error to be approximately one half the membrane thickness, 3.5 nm.

### Interpolation of Gaps Between Serial Sections

Both datasets are composed of a number of physical serial sections of neuropil, while spacing between the reconstructed virtual sections is 1.64 nm, the spacing between the physical sections is somewhat larger. When the physical sections are joined with the same 1.64 nm spacing, an obvious discontinuity in the shapes of the mitochondria and other subcellular structures is visible at the interface between one physical section and the next. This is especially apparent in the XZ and YZ projections of the volumes (Supplementary Figure 8A). As the alignment of the physical sections by SWiFT-IR was excellent, we surmised that the discontinuity was due to a gap (i.e. missing virtual sections) in the Z spacing. In order to estimate the proper Z spacing, the XYZ mode of IMOD was used to compile six cross sections from each dataset in the XZ or YZ planes (Supplementary Figure 8B). This allowed the degree of discontinuity to be assessed visually at six locations for each physical gap. The amount of Z spacing added to achieve continuity is an average for each of the six views. The average gap sizes for Dataset 1 and Dataset 2 are (15.6 ± 0.8) nm and (18.6 ± 1.6) nm, respectively. Gaps of these sizes are equivalent to approximately 10 and 11 missing virtual tomographic sections at a thickness of 1.64 nm. In order to create complete reconstructions, we inserted 10 blank virtual sections at each gap and additional contours, by manual interpolation across the gap, assisted by IMOD’s spherical and linear interpolation tools [21].

### Mesh Generation

The [CellBlender](https://mcell.org/) [3] modeling plug-in for Blender was used for 3D mesh generation and analysis. The program Contour Tiler [10], which is integrated with the [NeuropilTools module](https://github.com/mcellteam/neuropil_tools) of CellBlender, was employed to create triangulated surface meshes from the traced contours. The resulting surface meshes retain the exact shape and dimensions of each contour down to the sub-nanometer level. This was performed separately for the three membrane objects in each mitochondrion. A Blender “Boolean Difference Modifier” was subsequently applied to subtract the cristae membrane object from the inner boundary membrane object (Figure 1E-F). This created the cavities in the inner boundary membrane due to the cristae junctions and cristae membrane. In the operation the protrusions disappeared and the cristae junction curvature was preserved. Although no alterations have been made to the traces at this stage, the intrinsic human tracing errors make the mesh jagged and uneven. To improve the meshes, we used the Smooth and Normal Smooth mesh improvement tools in [GAMer 2](https://github.com/ctlee/gamer) [23]. The noise of the improved surface mesh is greatly reduced, and when overlayed on the original noisey mesh is seen to pass between the peaks and valleys of the noise of the original boundary traces, thus reducing the tracing error. Finally to verify if any compression artifact had occurred in the 3DEM dataset we traced and reconstructed synaptic vesicles. Analysis of the shapes of the vesicles revealed 23.9% shrinkage in the Z direction. We scaled the reconstructed meshes by a factor of 1.27 in the Z direction to obtain spherical vesicles of 40 nm diameter. This Z-scaling factor was then applied to the mitochondria meshes.

### Mesh Analysis

All mesh analysis and characterization were performed in Blender using Blender add-ons. Surface area, volume and genus were measured with Cellblender mesh analysis, and linear distances were computed with MeasureIT. Skeletons of the outer and inner membrane were generated with the mesh [skeletonization tool](https://doc.cgal.org/latest/Surface_mesh_skeletonization/index.html) of the CGal library [50], interfaced in Blender through the python API (this generated code is also provided). For this purpose meshes needed to be converted to OFF format with a Blender [OFF Add-on](https://github.com/alextsui05/blender-off-addon). The skeletons were further used to calculate the radius of the organelle at positions along its length, projecting points from the skeleton to the mesh, and calculating the radial distance to it (Supplementary Figure 9). Several points are used to measure the radius at each location in the skeleton, all with a distance smaller than 0.01 μm to the perpendicular plane set at the location, this is the source of errors in Supplementary Figure 9. With a similar approach we calculated the length of the organelles.

We estimated the diameter of the cristae junctions from the measurements of the area of the inner boundary membrane with and without cristae junctions (i.e. before and after the performed Boolean operation).

The average surface area taken up by CJs is: CJ area = (SA IBM closed - SA IBM)/# CJs. We further assume the cristea junctions area circular, and from this area we calculated the radius: CJ area = *πr*^2^.

Curvature analysis was carried out with GAMer 2 [24, 23] through its Blender interface, using the MDSB algorithm [30] for curvature calculation.

Idealized shapes were generated with Netgen [42]. All statistical analysis and plots were generated using Python 3.8, with the libraries NumPy [16], SciPy [52], Matplotlib [19] and Scikit-learn [34]. The K-means clustering algorithm implemented in Scikit-learn was used to cluster geometric features of the organelles. Given a sets of points (2 dimensional in our case), this algorithm partitions the points into k sets (k = 2 in our case) minimizing the sum of squares (i.e. the variance). Statistical significance was assessed using a non-parametric method, the two-sample Kolmogorov-Smirnov (KS) test.

## Acknowledgments

We would like to acknowledge support from Air Force Office of Scientific Research (AFOSR) Multidisciplinary University Research Initiative (MURI) FA9550-18-1-0051 to P.R., T.J.S, T.M.B., R.M, G.C.G. T.J.S.: NIH GM103712, NSF DBI-1707356 and NSF DBI-2014862. P.R: Office of Naval Research N00014-20-1-2469 G.C.G. was partially supported by Luxembourg National Research Fund in the frame of a PhD Grant No.9984574. C.T.L. was supported by a Hartwell Foundation postdoctoral fellowship. We would like to thank Kelly Brockmeyer, Aranza S.M. Lopez, Andrea S. Jacinto, Justin Oshiro, Andrew Nguyen, and Meagan P. Rowan for helping with image segmentation and preliminary work. We also thank Prof. Kristen Harris for discussion and suggestions. The funders had no role in study design, data collection and analysis, decision to publish, or preparation of the manuscript.

## Supplementary Material

**Table 4:** Quantification of the geometric properties of the outer membrane: volume, surface area, length and average radius.

| Mito reference # | OM Volume ($\mu\text{m}^3$) | OM surface area ($\mu\text{m}^2$) | Length ($\mu\text{m}$) | Average radius ($\mu\text{m}$) |
| --- | --- | --- | --- | --- |
| 1 | 0.0360 | 0.6325 | 0.797 | 0.132 |
| 9 | 0.0102 | 0.2343 | 0.307 | 0.128 |
| 10 | 0.0140 | 0.3577 | 0.532 | 0.1 |
| 11 | 0.0062 | 0.1788 | 0.350 | 0.08 |
| 12 | 0.0124 | 0.2655 | 0.332 | 0.134 |
| 19 | 0.0205 | 0.397 | 0.475 | 0.131 |
| 5 | 0.0459 | 0.7968 | 0.981 | 0.129 |
| 4 | 0.0287 | 0.6 | 0.895 | 0.108 |
| 6 | 0.0123 | 0.2816 | 0.445 | 0.11 |
| 7 | 0.0121 | 0.3372 | 0.674 | 0.082 |
| 8 | 0.0152 | 0.351 | 0.590 | 0.092 |
| 14 | 0.0092 | 0.2558 | 0.434 | 0.087 |

**Figure 8:**
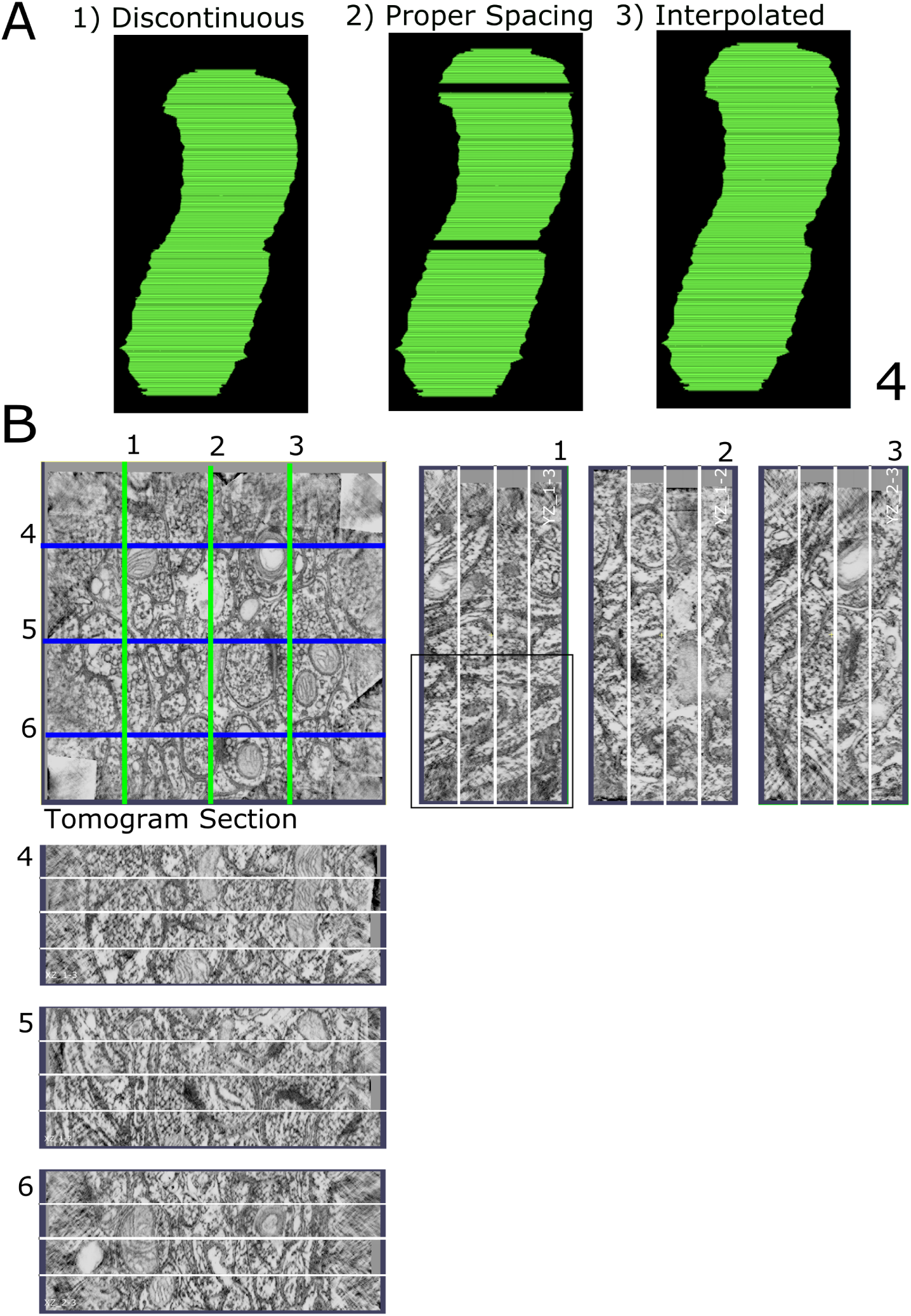
Gap size determination and contour interpolation. A) Shown in A1 is a mitochondrion with uniform spacing of 1.64 nm between all tomographs. In A2, appropriate spacing of 18 nm has been added to account for gaps between physical sections. A3 shows the gaps filled in with new contours using manual and automatic methods in IMOD. B) Shown here are six side views compiled from a set of tomograms exemplified by the top left image. The space seen at the junction between images was increased until the membranes could be connected by a smooth, continuous curve rather than a jagged line. The length of these spaces was averaged across the 6 views for each of the 3 gaps. The average of 18.5 nm was translated into 11 sections at the uniform 1.64 nm thickness.

**Figure 9:**
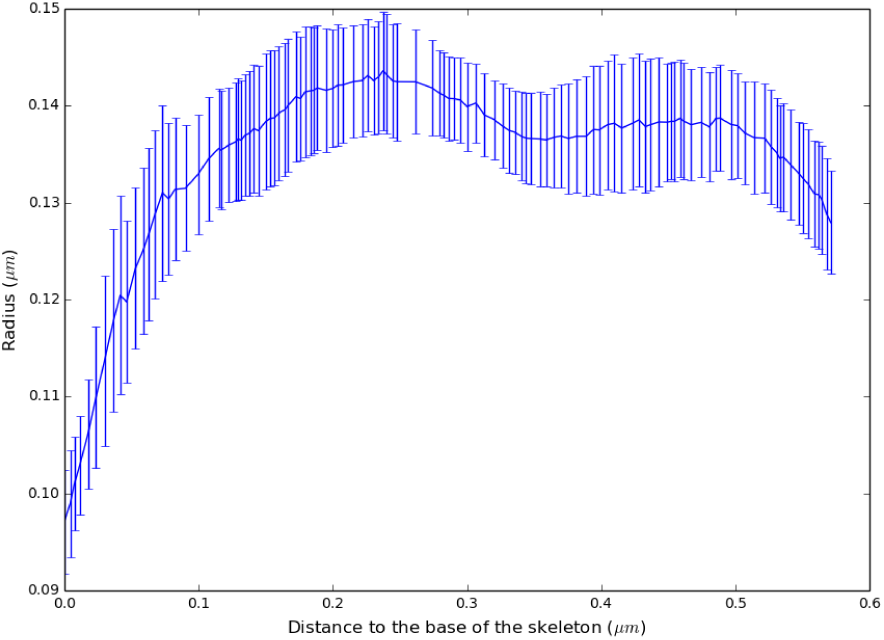
Representative figure of the radius along the skeleton for the OM of one organelle.

**Figure 10:**
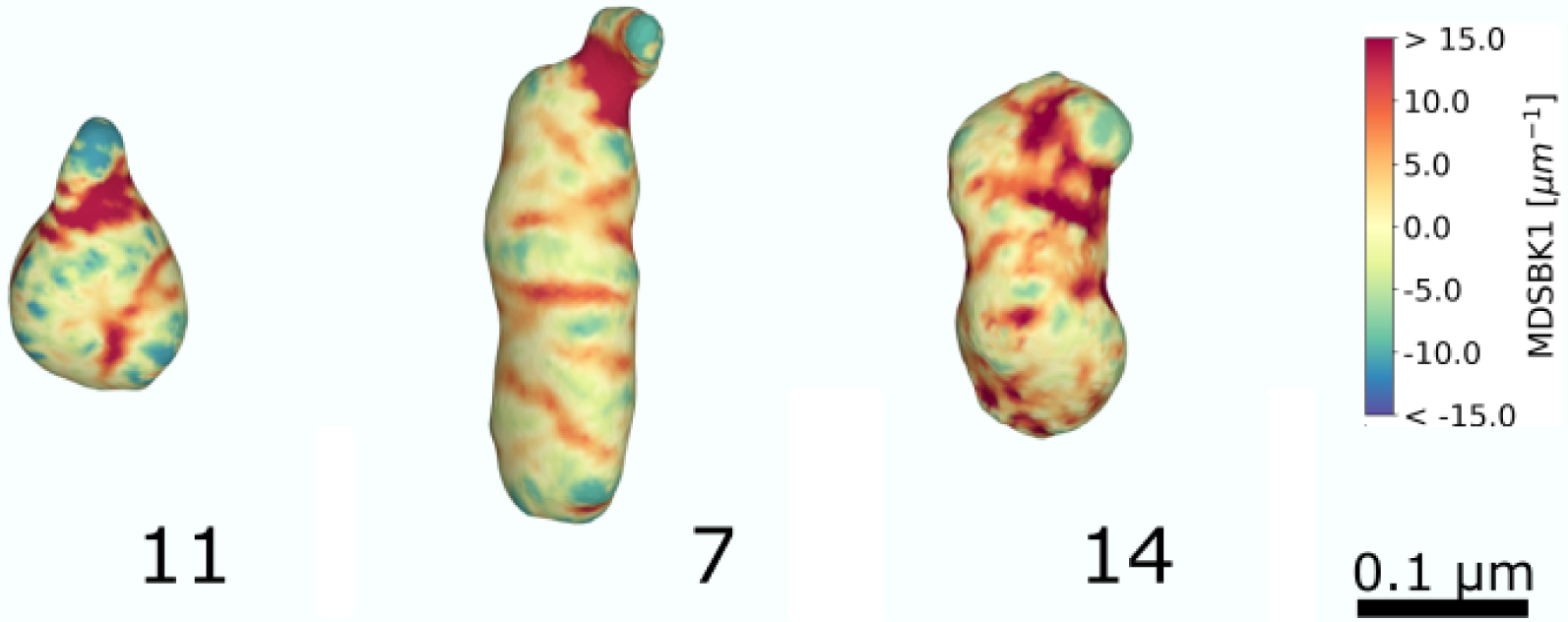
Curvature estimation for the OM with protrusions.

**Figure 11:**
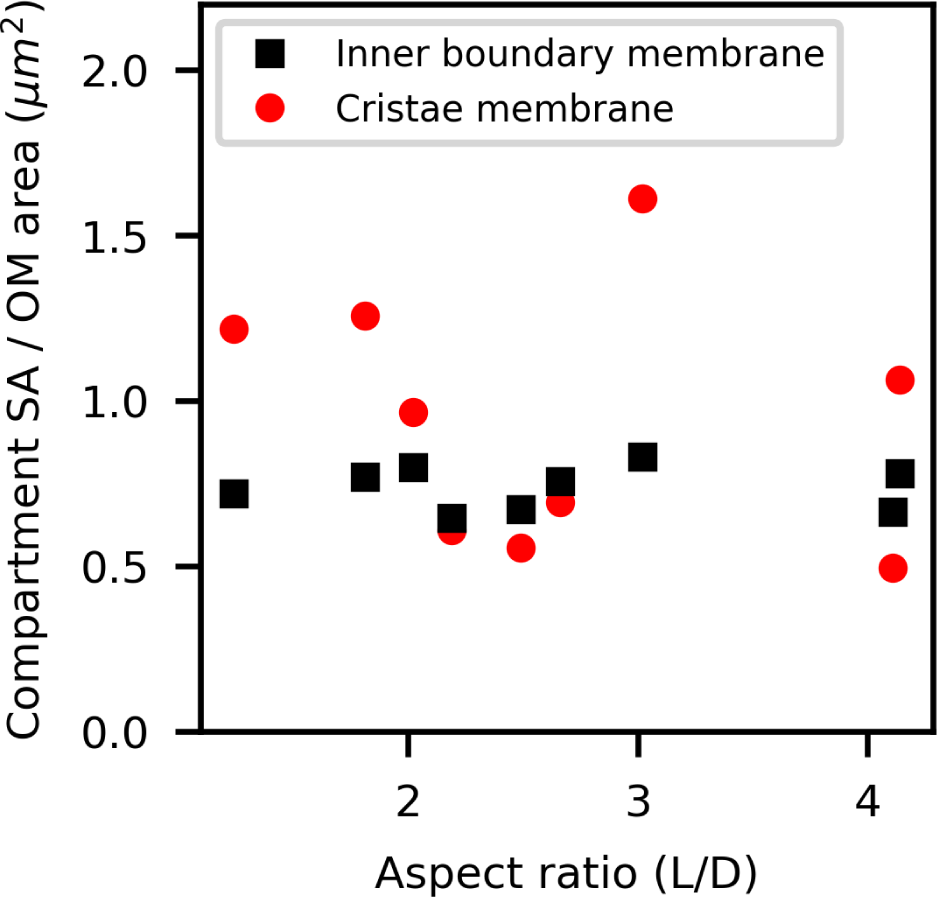
Ratio of the surface area of the inner boundary membrane and cristae membrane to the outer membrane surface area.

**Figure 12:**
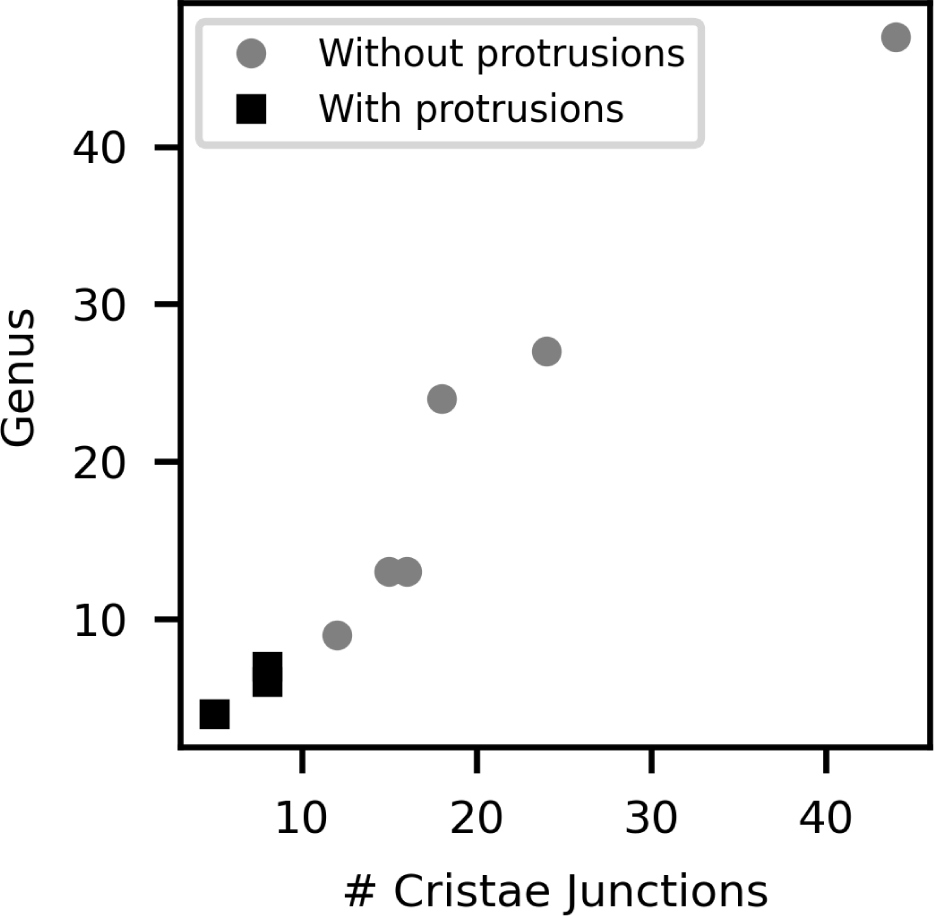
The genus of the inner membrane is significantly smaller for mitochondria with protrusions. This quantity is highly correlated with the number of cristae junctions.

**Figure 13:**
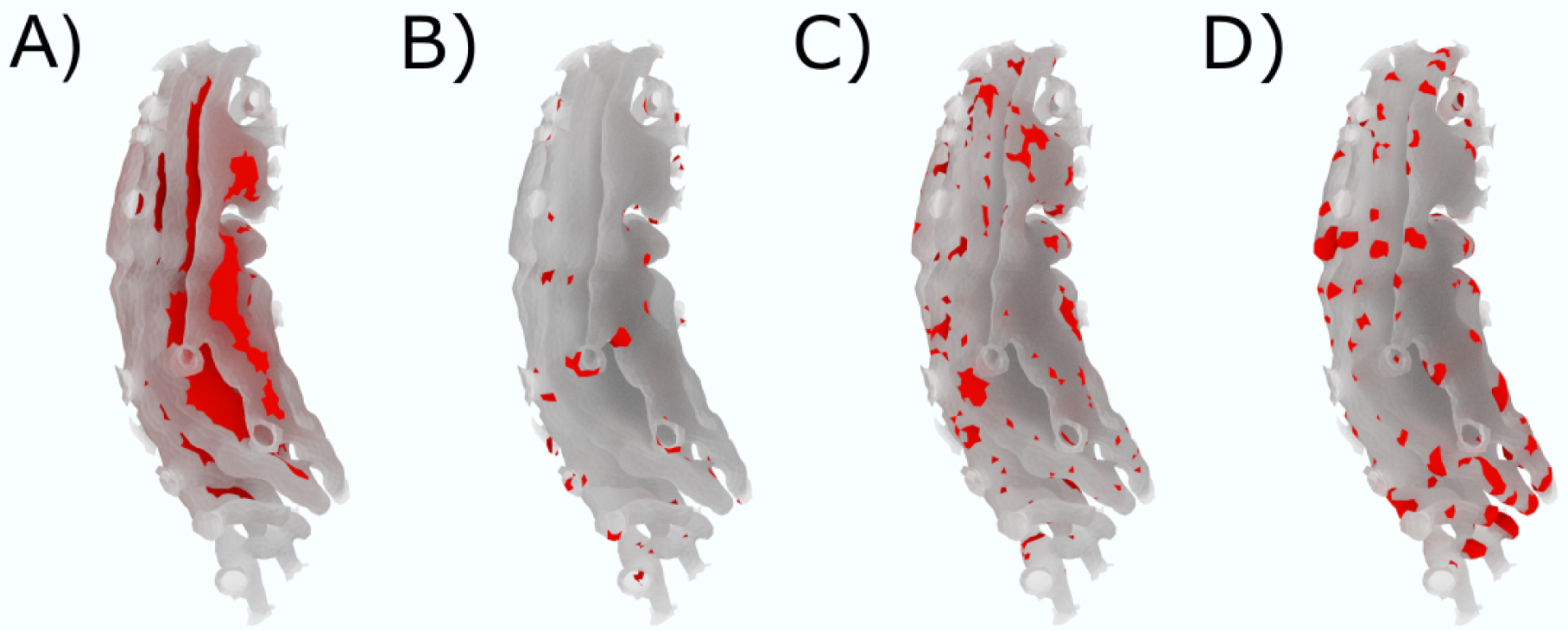
Structural motif representation. (A) Flat-sheets (B) Saddle-like structures (C) Semi-cylindrical shapes (D) Semi-spherical shapes.

**Figure 14:**
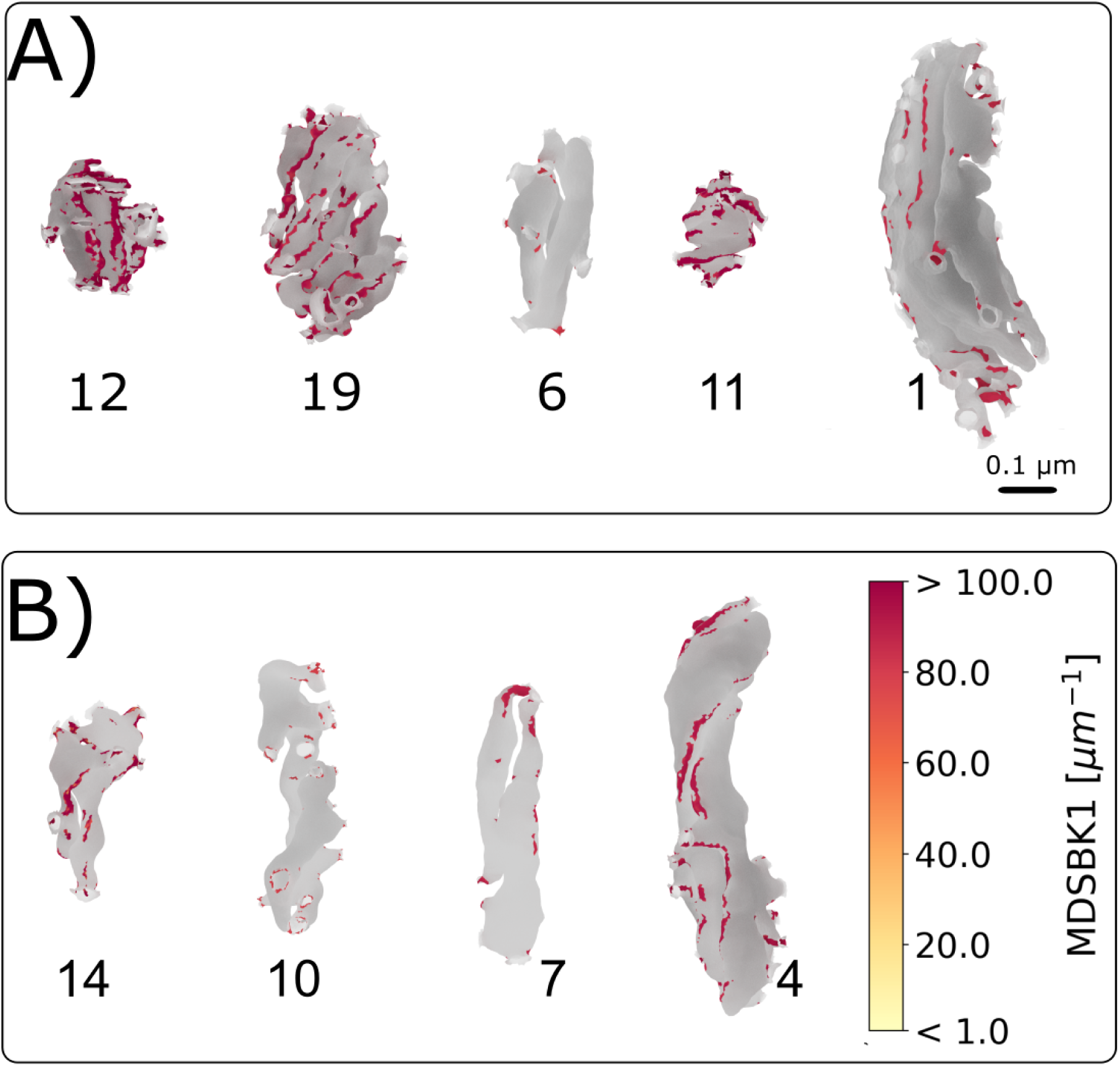
High curvature regions of the inner mitochondrial membrane. A-B) Only vertices of the cristae membrane with the first principal curvature k1 greater than 80 μm^−1^ are shown in color according to the color bar. Remaining vertices are in grey. (A) Globular mitochondria population and (B) Elongated mitochondria population.

## Notes

### Competing Interest Statement

The authors have declared no competing interest.

## References

[1] Haleh Alimohamadi and Padmini Rangamani. “Modeling membrane curvature generation due to membrane-protein interactions”. In: Biomolecules 8.4 (2018), p. 120. url: https://doi.org/10.3390/biom8040120.

[2] David Attwell and Simon B Laughlin. “An energy budget for signaling in the grey matter of the brain”. In: Journal of Cerebral Blood Flow & Metabolism 21.10 (2001), pp. 1133–1145. url: https://doi.org/10.1097/00004647-200110000-00001.

[3] Thomas M. Bartol et al. “Computational reconstitution of spine calcium transients from individual proteins”. In: Frontiers in Synaptic Neuroscience 7 (2015), p. 17. issn: 1663-3563. url: https://doi.org/10.3389/fnsyn.2015.00017.

[4] Miriam Bell et al. “Dendritic spine geometry and spine apparatus organization govern the spatiotemporal dynamics of calcium”. In: Journal of General Physiology 151.8 (2019), pp. 1017–1034. url: https://doi.org/10.1085/jgp.201812261.

[5] Morgan Chabanon and Padmini Rangamani. “Gaussian curvature directs the distribution of spontaneous curvature on bilayer membrane necks”. In: Soft matter 14.12 (2018), pp. 2281–2294. url: https://doi.org/10.1039/c8sm00035b.

[6] Sara Cogliati et al. “Mitochondrial cristae shape determines respiratory chain supercomplexes assembly and respiratory efficiency”. In: Cell 155.1 (2013), pp. 160–171. url: https://doi.org/10.1016/j.cell.2013.08.032.

[7] Csaba Cserép et al. “Mitochondrial ultrastructure is coupled to synaptic performance at axonal release sites”. In: Eneuro 5.1 (2018). url: https://doi.org/10.1523/ENEURO.0390-17.2018.

[8] Andrea Cugno et al. “Geometric principles of second messenger dynamics in dendritic spines”. In: Scientific reports 9.1 (2019), pp. 1–18. url: https://doi.org/10.1038/s41598-019-48028-0.

[9] Michael J Devine and Josef T Kittler. “Mitochondria at the neuronal presynapse in health and disease”. In: Nature Reviews Neuroscience 19.2 (2018), p. 63. url: https://doi.org/10.1038/nrn.2017.170.

[10] John Edwards et al. “VolRoverN: enhancing surface and volumetric reconstruction for realistic dynamical simulation of cellular and subcellular function”. In: Neuroinformatics 12.2 (2014), pp. 277–289. url: https://doi.org/10.1007/s12021-013-9205-2.

[11] John C Fiala. “Reconstruct: a free editor for serial section microscopy”. In: Journal of microscopy 218.1 (2005), pp. 52–61. url: https://doi.org/10.1111/j.1365-2818.2005.01466.x.

[12] Terrence G Frey and Carmen A Mannella. “The internal structure of mitochondria”. In: Trends in biochemical sciences 25.7 (2000), pp. 319–324. url: https://doi.org/10.1016/s0968-0004(00)01609-1.

[13] Terrence G Frey, Guy A Perkins, and Mark H Ellisman. “Electron tomography of membrane-bound cellular organelles”. In: Annu. Rev. Biophys. Biomol. Struct. 35 (2006), pp. 199–224. url: https://doi.org/10.1146/annurev.biophys.35.040405.102039.

[14] Guadalupe C Garcia et al. “Mitochondrial morphology provides a mechanism for energy buffering at synapses”. In: Scientific reports 9 (2019). url: https://doi.org/10.1038/s41598-019-54159-1.

[15] Marta Giacomello et al. “The cell biology of mitochondrial membrane dynamics”. In: Nature Reviews Molecular Cell Biology (2020), pp. 1–21. url: https://doi.org/10.1038/s41580-020-0210-7.

[16] Charles R. Harris et al. “Array programming with NumPy”. In: Nature 585 (2020), pp. 357–362. doi: 10.1038/s41586-020-2649-2.

[17] Julia J Harris, Renaud Jolivet, and David Attwell. “Synaptic energy use and supply”. In: Neuron 75.5 (2012), pp. 762–777. url: https://doi.org/10.1016/j.neuron.2012.08.019.

[18] Wolfgang Helfrich. “Elastic properties of lipid bilayers: theory and possible experiments”. In: Zeitschrift für Naturforschung C 28.11-12 (1973), pp. 693–703. url: https://doi.org/10.1515/znc-1973-11-1209.

[19] J. D. Hunter. “Matplotlib: A 2D Graphics Environment”. In: Computing in Science Engineering 9.3 (2007), pp. 90–95. doi: 10.1109/MCSE.2007.55.

[20] Arun Kumar Kondadi et al. “Cristae undergo continuous cycles of membrane remodelling in a MICOS-dependent manner”. In: EMBO reports 21.3 (2020), e49776. url: https://doi.org/10.15252/embr.201949776.

[21] James R Kremer, David N Mastronarde, and J Richard McIntosh. “Computer visualization of three-dimensional image data using IMOD”. In: Journal of structural biology 116.1 (1996), pp. 71–76. url: https://doi.org/10.1006/jsbi.1996.0013.

[22] Erwin Kreyszig. Introduction to differential geometry and Riemannian geometry. University of Toronto Press, 1968. url: https://doi.org/10.3138/9781487589448.

[23] Christopher T. Lee et al. “3D Mesh Processing Using GAMer 2 to Enable Reaction-Diffusion Simulations in Realistic Cellular Geometries”. In: PLOS Computational Biology 16.4 (Apr. 2020), e1007756. issn: 1553-7358. url: https://doi.org/10.1371/journal.pcbi.1007756.

[24] Christopher T. Lee et al. “An Open-Source Mesh Generation Platform for Biophysical Modeling Using Realistic Cellular Geometries”. In: Biophysical Journal 118.5 (Mar. 2020), pp. 1003–1008. issn: 0006-3495. url: https://doi.org/10.1016/j.bpj.2019.11.3400.

[25] A Leung et al. “Deciphering the postsynaptic calcium-mediated energy homeostasis through mitochondria-endoplasmic reticulum contact sites using systems modeling”. In: bioRxiv (2020). url: https://doi.org/10.1101/2020.09.12.294827.

[26] Gaurang Mahajan and Suhita Nadkarni. “Intracellular calcium stores mediate metaplasticity at hippocampal dendritic spines”. In: The Journal of Physiology 597.13 (2019), pp. 3473–3502. url: https://doi.org/10.1113/JP277726.

[27] Carmen A Mannella. Introduction: our changing views of mitochondria. 2000. url: https://doi.org/10.1023/a:1005562109678.

[28] Wallace F Marshall. “Differential geometry meets the cell”. In: Cell 154.2 (2013), pp. 265–266. url: https://doi.org/10.1016/j.cell.2013.06.032.

[29] Harvey T McMahon and Jennifer L Gallop. “Membrane curvature and mechanisms of dynamic cell membrane remodelling”. In: Nature 438.7068 (2005), pp. 590–596. url: https://doi.org/10.1038/nature04396.

[30] Mark Meyer et al. “Discrete differential-geometry operators for triangulated 2-manifolds”. In: Visualization and mathematics III. Springer, 2003, pp. 35–57.

[31] Thomas Misgeld and Thomas L Schwarz. “Mitostasis in neurons: maintaining mitochondria in an extended cellular architecture”. In: Neuron 96.3 (2017), pp. 651–666. url: https://doi.org/10.1016/j.neuron.2017.09.055.

[32] Nelli Mnatsakanyan and Elizabeth Ann Jonas. “The new role of F1Fo ATP synthase in mitochondria-mediated neurodegeneration and neuroprotection”. In: Experimental Neurology (2020), p. 113400. url: https://doi.org/10.1016/j.expneurol.2020.113400.

[33] Patrick Paumard et al. “The ATP synthase is involved in generating mitochondrial cristae morphology”. In: The EMBO journal 21.3 (2002), pp. 221–230. url: https://doi.org/10.1093/emboj/21.3.221.

[34] Fabian Pedregosa et al. “Scikit-learn: Machine Learning in Python”. In: Journal of Machine Learning Research 12.85 (2011), pp. 2825–2830. url: http://jmlr.org/papers/v12/pedregosa11a.html.

[35] G Perkins et al. “Electron tomography of neuronal mitochondria: three-dimensional structure and organization of cristae and membrane contacts”. In: Journal of structural biology 119.3 (1997), pp. 260–272. url: https://doi.org/10.1006/jsbi.1997.3885.

[36] Guy Perkins, Dakota R. Jackson, and George A. Spirou. “Resolving presynaptic structure by electron tomography”. In: Synapse 69 (2015), pp. 268–282. url: https://doi.org/10.1002/syn.21813.

[37] Guy Perkins et al. “Altered outer hair cell mitochondrial and subsurface cisternae connectomics are candidate mechanisms for hearing loss in mice”. In: Journal of Neuroscience 40.44 (2020), pp. 8556–8572. url: https://doi.org/10.1523/JNEUROSCI.2901-19.2020.

[38] Guy A Perkins, Mark H Ellisman, and Donald A Fox. “Three-dimensional analysis of mouse rod and cone mitochondrial cristae architecture: bioenergetic and functional implications”. In: Mol Vis 9 (2003), pp. 60–73. url: http://europepmc.org/abstract/MED/12632036.

[39] Guy A Perkins et al. “Membrane architecture of mitochondria in neurons of the central nervous system”. In: Journal of Neuroscience Research 66.5 (2001), pp. 857–865. url: https://doi.org/10.1002/jnr.10050.

[40] Guy A Perkins et al. “The micro-architecture of mitochondria at active zones: electron tomography reveals novel anchoring scaffolds and cristae structured for high-rate metabolism”. In: Journal of Neuroscience 30.3 (2010), pp. 1015–1026. url: https://doi.org/10.1523/JNEUROSCI.1517-09.2010.

[41] Sébastien Phan et al. “3D reconstruction of biological structures: automated procedures for alignment and reconstruction of multiple tilt series in electron tomography”. In: Advanced Structural and Chemical Imaging 2.1 (2016), p. 8. url: https://doi.org/10.1186/s40679-016-0021-2.

[42] Joachim Schöberl. “NETGEN An advancing front 2D/3D-mesh generator based on abstract rules”. In: Computing and visualization in science 1.1 (1997), pp. 41–52. url: https://doi.org/10.1007/s007910050004.

[43] Stephanie E Siegmund et al. “Three-Dimensional Analysis of Mitochondrial Crista Ultrastructure in a Patient with Leigh Syndrome by In Situ Cryoelectron Tomography”. In: iScience 6 (2018), pp. 83–91. url: https://doi.org/10.1016/j.isci.2018.07.014.

[44] Nishant Singh et al. “Presynaptic endoplasmic reticulum regulates short-term plasticity in hippocampal synapses”. In: Communications Biology 4.1 (2021), pp. 1–13. url: https://doi.org/10.1038/s42003-021-01761-7.

[45] Gina E. Sosinsky et al. “The combination of chemical fixation procedures with high pressure freezing and freeze substitution preserves highly labile tissue ultrastructure for electron tomography applications”. In: Journal of Structural Biology 161.3 (2008). The 4th International Conference on Electron Tomography, pp. 359–371. issn: 1047-8477. url: https://doi.org/10.1016/j.jsb.2007.09.002.

[46] Till Stephan et al. “MICOS assembly controls mitochondrial inner membrane remodeling and crista junction redistribution to mediate cristae formation”. In: The EMBO journal 39.14 (2020), e104105. url: https://doi.org/10.15252/embj.2019104105.

[47] Stefan Stoldt et al. “Mic60 exhibits a coordinated clustered distribution along and across yeast and mammalian mitochondria”. In: Proceedings of the National Academy of Sciences 116.20 (2019), pp. 9853–9858. url: https://doi.org/10.1073/pnas.1820364116.

[48] Mike Strauss et al. “Dimer ribbons of ATP synthase shape the inner mitochondrial membrane”. In: The EMBO journal 27.7 (2008), pp. 1154–1160. url: https://doi.org/10.1038/emboj.2008.35.

[49] Dirk Jan Struik. Lectures on classical differential geometry. Courier Corporation, 1961.

[50] Andrea Tagliasacchi et al. “Mean curvature skeletons”. In: Computer Graphics Forum. Vol. 31. 5. Wiley Online Library. 2012, pp. 1735–1744. url: https://doi.org/10.1111/j.1467-8659.2012.03178.x.

[51] Mark Terasaki et al. “Stacked endoplasmic reticulum sheets are connected by helicoidal membrane motifs”. In: Cell 154.2 (2013), pp. 285–296. url: https://doi.org/10.1016/j.cell.2013.06.031.

[52] Pauli Virtanen et al. “SciPy 1.0: Fundamental Algorithms for Scientific Computing in Python”. In: Nature Methods 17 (2020), pp. 261–272. doi: 10.1038/s41592-019-0686-2.

[53] Eric W Weisstein. Catenoid From MathWorld–A Wolfram Web Resource. 2013. url: https://mathworld.wolfram.com/Catenoid.html.

[54] Eric W Weisstein. Cylinder From MathWorld–A Wolfram Web Resource. url: https://mathworld.wolfram.com/Cylinder.html.

[55] Eric W Weisstein. Hyperbolic Paraboloid From MathWorld–A Wolfram Web Resource. 2014. url: http://mathworld.wolfram.com/HyperbolicParaboloid.html..

[56] Eric W Weisstein. Sphere From MathWorld–A Wolfram Web Resource. 2014. url: https://mathworld.wolfram.com/Sphere.html.

[57] Arthur W Wetzel et al. “Registering large volume serial-section electron microscopy image sets for neural circuit reconstruction using FFT signal whitening”. In: 2016 IEEE Applied Imagery Pattern Recognition Workshop (AIPR). IEEE. 2016, pp. 1–10. url: https://doi.org/10.1109/AIPR.2016.8010595.

[58] Joshua Zimmerberg and Michael M Kozlov. “How proteins produce cellular membrane curvature”. In: Nature reviews Molecular cell biology 7.1 (2006), pp. 9–19. url: https://doi.org/10.1038/nrm1784.

